# A comprehensive study of Phospholipid fatty acid rearrangements in the early onset of the metabolic syndrome: correlations to organ dysfunction

**DOI:** 10.1101/2019.12.13.875096

**Authors:** Amélie Bacle, Linette Kadri, Spiro Khoury, Romain Ferru-Clément, Jean-François Faivre, Christian Cognard, Jocelyn Bescond, Amandine Krzesiak, Hugo Contzler, Nathalie Delpech, Jenny Colas, Clarisse Vandebrouck, Stéphane Sébille, Thierry Ferreira

## Abstract

The balance within phospholipids (PL) between Saturated Fatty Acids (SFA) and mono- or poly-Unsaturated Fatty Acids (UFA), is known to regulate the biophysical properties of cellular membranes. As a consequence, perturbating this balance alters crucial cellular processes in many cell types, such as vesicular budding and the trafficking/function of membrane-anchored proteins. The worldwide spreading of the Western-diet, which is specifically enriched in saturated fats, has been clearly correlated with the emergence of a complex syndrome, known as the Metabolic Syndrome (MetS), which is defined as a cluster of risk factors for cardiovascular diseases, type 2 diabetes and hepatic steatosis. However, no clear correlations between diet-induced fatty acid redistribution within cellular PL, the severity/chronology of the symptoms associated to MetS and the function of the targeted organs, particularly in the early onset of the disease, have been established. In an attempt to fill this gap, we analyzed in the present study PL remodeling in rats exposed during 15 weeks to a High Fat/High Fructose diet (HFHF) in several organs, including known MetS targets. We show that fatty acids from the diet can distribute within PL in a very selective way, with PhosphatidylCholine being the preferred sink for this distribution. Moreover, in the HFHF rat model, most organs are protected from this redistribution, at least during the early onset of MetS, at the exception of the liver and skeletal muscles. Interestingly, such a redistribution correlates with clear-cut alterations in the function of these organs.

## 1. Introduction

The first observations concerning the involvement of obesity and dyslipidemia in the occurrence of metabolic disorders, including type 2 diabetes, fatty liver and cardiovascular diseases, go back to the late 1960’s. Since then, the prevalence of this metabolic syndrome has been clearly correlated to the worldwide spreading of the Western-diet which is excessively rich in sugar and saturated fat. In obese individuals, the incidence of the metabolic syndrome is associated with high plasma levels of Non Esterified Fatty Acids (NEFA) and more specifically of long-chain saturated fatty acids (SFA) (*1*). A prime example of long chain SFA is Palmitate (bearing 16 carbons and no double-bond in the acyl chain: 16:0), the main component of Palm oil. When the storage capacity of the adipose tissue is exceeded, NEFA tend to accumulate into cells not suited for lipid storage, among which muscle cells, hepatocytes and pancreatic β-cells are prime examples (*1*). As a corollary, it is now widely accepted that fatty acid imbalances are directly involved in the promotion of insulin resistance, non-alcoholic steatohepatitis (NASH), impaired glucose tolerance and systemic inflammation (*2*).

Phospholipids (PL), which bear two fatty acid chains, are the main components of cellular membranes. In mammalian cells, PhosphatidylCholine (PC) is the most abundant PL (*3*). Ethanolamine (Etn)-containing PL species are the second most abundant phospholipids, in which PhosphatidylEthanolamine (PE), a diacyl glycerophospholipid, and ethanolamine plasmalogen (PE(P)), an alkenylacylglycerophospholipid, are the main constituents (*4, 5*). These species, and to a lesser extent, PhosphatidylInositol (PI) and PhosphatidylSerine (PS), are the most abundant lipid classes whatever the organ considered, constituting 75 % of all lipid species in the heart of rats, and 79 % in the liver, as examples (*3*). Maintaining the equilibrium between SFA, Mono- and Poly-Unsaturated Fatty Acids (UFA) within membrane PL is crucial to sustain the optimal membrane biophysical properties, compatible with selective organelle based processes, including vesicular budding or membrane-protein trafficking and function (*6*). As a corrolary, impaired balances within SFA and UFA have been demonstrated to result in dramatic cellular dysfunctions in cells relevant of the metabolic syndrome (for review, see (*7*)). As examples, altered insulin secretion in pancreatic β-cells or impairment in Glucose disposal in muscle and liver cells have been reported in response to SFA overload (*7*). This process, which is referred to as lipotoxicity, ultimately leads to cell death by apoptosis in all the cellular systems tested (*7*).

Many animal models of the metabolic syndrome, either genetic or diet-induced, have been described so far (*8*). Recently, Lozano *et al.* reported an elegant rat model based on an high-fructose and high-fat diet (HFHF; (*9*)). These authors could demonstrate that the combination of high fat and high carbohydrate induced type 2 diabetes with widespread tissues effects. The phenotype increased gradually from two month to eight months following the shift to HFHF, with insulin-resistance and hepatic disorders increasing progressively to reach a maximum at this latter time point. Since with the increased consumption of sugar-rich and fatty-products, and the increase in preference for such products, metabolic disorders are becoming more common at a younger age in humans, this model appears to be very appealing to evaluate the chronology of the impacts of the diet, particularly during the early onset of the metabolic syndrome.

In this context, the present study aimed at evaluating the distribution of fatty acids coming from the diet within cellular PL within various organs in the HFHF model. We focused in this work on the early stages of the phenotype set-up. Moreover, the function of the most impacted organs, in terms of fatty acid distribution, was also evaluated.

## 2. Materials and Methods

### 2.1 Animals

The present study was approved by the Comité d’Ethique et d’Expérimentation Animale (COMETHEA) and the French Ministère de l’Enseignement Supérieur, de la Recherche et de l’Innovation (authorization n°2016071215184098). The protocols were designed according to the Guiding Principles in the Care and Use of Animals approved by the Council of the American Physiological Society and were in adherence with the Guide for the Care and Use of Laboratory Animals published by the US National Institutes of Health (NIH Publication no. 85-23, revised 1996) and according to the European Parliament Directive 2010/63 EU.

We recapitulated the model developed by Lozano *et al.* (*9*). Twenty eight male eight week old Wistar rats (275-299 g), supplied by Envigo (Gannat, France), were housed in a temperature-controlled room, in a 12-h-light/dark cycle environment with *ad libitum* access to water and food. After one week of acclimatization, the rats were randomly divided into two groups of 14 rats each. The first group had free access to a standard diet (CTL) from MUCEDOLA (Settimo Milanese, Italy), with the following macronutrient composition: 3.0 % fat, 18.5 % protein, 46 % carbohydrate, 6 % fibre, and 7 % ash (minerals). The second group “High Fat High Fructose” (HFHF) received a purified laboratory hypercaloric rodent diet “WESTERN RD” (SDS, Special Diets Services, Saint Gratien, France) containing 21.4 % fat, 17.5 % protein, 50 % carbohydrate, 3.5 % fibre, and 4.1 % ash, and additional 25 % of fructose (Sigma-Aldrich, Saint-Louis, Missouri, USA) in water. Fatty acid distribution within the fat fraction of both diets is displayed in Supplementary Table 1. In “WESTERN RD”, among fatty acids, the mono-unsaturated fatty acid Oleate and the saturated fatty acid Palmitate were the most represented and were found in equal amounts, contributing to 60 % of total fatty acids within the fat fraction. During the longitudinal observation, body weight was measured each week and plasma glucose, plasma Triglycerides and plasma Cholesterol levels were determined every 2 weeks at the same time (2 PM) under random fed conditions. A final experiment was performed 15 weeks after the initiation of the different diets and plasma glucose, insulin, cholesterol, triglyceride and NEFA levels were measured after a three hour fasting period to allow gastric emptying. All rats were sacrificed 16 weeks after starting administration of each diet for futher lipidomic profiling and *ex-vivo* experiments.

### 2.2 Biochemical plasma analysis

Blood glucose levels were measured using an automatic glucose monitor (One Touch vita - LifeScan Inc., Milpitas, California, USA). Plasma triglyceride and total Cholesterol concentrations were determined using commercially available colorimetric kits (Sobioda, Montbonnot-Saint-Martin, France). Plasma free fatty acids were quantified by a colorimetric NEFA kit (Wako Chemicals, Osaka, Japan). For lipoprotein characterization, plasma samples were collected and subjected to fractionation by FPLC (ÄKTA pure chromatography system - GE Healthcare Life Sciences, Chicago, Illinois, USA). Cholesterol and Triglyceride concentrations in each fraction were measured using commercially available colorimetric kits (Sobioda, Montbonnot-Saint-Martin, France).

### 2.3 Lipid Extraction, Phospholipid Purification and Mass Spectrometry Analyses

After rats were anesthetized, the organs were quickly removed and put on ice surface. These organs were cut into small pieces (1-2 mm^3^) and dipped in liquid nitrogen. The frozen pieces were introduced into cryotubes before immersion in liquid nitrogen for storage at −80°C.

Lipids were extracted from each individual sample, according to the following procedure. Each frozen sample was first submitted to three rounds of grinding using a Precellys Evolution homogenizer (Bertin Technologies, Montigny-le-Bretonneux, France) and resuspended into 1 ml of water before tranfer into glass tubes containing 500 μL of glass beads (diameter 0.3–0.4 mm; Sigma-Aldrich, Saint-Louis, Missouri, USA). Lipids were extracted using chloroform/methanol (2:1, v/v) and shaking with an orbital shaker (IKAH VXR basic VibraxH - Sigma-Aldrich, Saint-Louis, Missouri, USA) at 1500 rpm during at least 1 h, as already described elsewhere (*10*). The final organic phase was evaporated and dissolved in 100 μL dichloromethane for purification of Phospholipids (PL) on a silica column (Bond ELUT-SI - Agilent Technologies, Santa Clara, California, USA). Lipid samples were loaded on the top of the column. Non-polar lipids were eluted by addition of 2 mL dichloromethane and glycolipids with 3 mL acetone. PLs were then eluted by 2 mL chloroform/methanol/H_2_O (50:45:5, v/v/v).

PL analysis in Mass Spectrometry (MS) was performed by a direct infusion of purified lipid extracts on a Synapt G2 HDMS (Waters Corporation, Milford, Massachusetts, USA) equipped with an Electrospray Ionization Source (ESI). The mass spectrum of each sample was acquired in the profile mode over 1 min. The scan range for PL analysis was from 500 to 1200 m/z. PS, PI, PE and its Plasmalogens (PE(P)) species were analyzed in negative ion mode after the addition of 0.1% (v/v) triethylamine (Supplementary Fig. 1A). PC species were analyzed in positive ion mode after the addition of 0.1% (v/v) formic acid as already described (*10*) (Supplementary Fig. 2A). Identification of the various PL species was based on their exact mass using the ALEX pipeline (*11*), and on MS/MS fragmentation for structural confirmation of the polar head (PL class) and the determincation of the fatty acid composition. MS/MS experiments of PI, PS, PE and PE(P) were performed by collision-induced-dissociation (CID) in the negative ion mode. Examples of obtained MS/MS spectra in the negative ion mode concerning some PLs are presented in the Supplementary Fig. 1B-D, which shows characteristic fragment ions allowing the identification of PL structures. MS/MS experiments in negative ion mode also allowed the identification of fatty acid chains in PC species as shown in Supplementary Fig. 2C. MS/MS spectrum in positive ion mode led to the identification of the polar head of PC class (characteristic and prominent fragment for all PC species with m/z 184, as shown in Supplementary Fig. 2B).

Compelementary experiments were also conducted on muscle and liver samples on non-purified lipid extracts in positive ion mode with a scan range from 300 to 1200 m/z to detect the neutral lipid species like Cholesterol, Diglycerides (DG) and Triglycerides (TG) (Supplementary Fig. 12). The structure of these compounds was confirmed based on the exact mass using the ALEX pipeline (*11*) and on MS/MS fragmentation of main species. All spectra were recorded with the help of MassLynx software (Version 4.1, Waters). Data processing of MS/MS spectra in this work was carried out with Biovia Draw 19.1© and MassLynx© software, with the help of Lipid Maps Lipidomics Gatway® (*https://www.lipidmaps.org/*)

### 2.4 Exercice

To determine the diet feeding’s effect on the functionnal capacity of rats, a maximal exercise test was used (*12*). In this test, 15 rats (CTL and HFHF groups, 7 and 8 per groups respectively) ran on a treadmill (Exer3/6 Treadmill - Columbus Instruments, Columbus, Ohio, USA) during five minutes at 13 m.min^-1^ at a grade of 10 degrees, and the speed was increased by 3.6 m.min^-1^ every two minutes until the animals were exhausted. At this time, the speed measured was their Maximum Running Speed (MRS). The week before the first MRS test, a treadmill habituation session was performed.

### 2.5 Muscles preparation and contraction measurement

EDL and Soleus muscles were carefully dissected with tendons intact on both ends and then vertically tied between a fixed hook at the bottom of the water jacketed 100 mL chamber (EmkaBATH2 - Emka Electronique, Noyant-la-Gravoyère, France) and the force transducer (MLTF500ST - ADInstruments, Dunedin, New Zealand) by means of cotton threads. Before experiments, muscles were maintained for 10 min into the 25°C physiological solution chamber, under oxygenated conditions (95% O_2_ and 5% CO_2_). The physiological solution (Krebs solution) contained (in mmol/L.): 120 NaCl, 5 KCl, 2 CaCl_2_, 1 MgCl2, 1 NaH_2_PO_4_, 25 NaHCO_3_, and 11 glucose) and pH was 7.4. The isometric tension was recorded by means of the transducer through a module (PowerLab 2/26 - ADInstruments, Dunedin, New Zealand) driven by the LabChart7 software (ADInstruments, Dunedin, New Zealand). Electrical external field stimulation was delivered through a constant current stimulator (STM4 - Bionic Instruments, Grenoble, France) and a pair of platinum electrodes (Radnoti, Terenure, Ireland) flanking both sides of the isolated muscle. Optimum stimulation conditions and muscle length were established in the course of preliminary experiments. In our device setup, supramaximal stimulation amplitude proved to be 200 mA for 1 ms, and the optimum length achieved for a resting pre-tension of 2 g.

The force-frequency relation was achieved by increasing step by step the frequency of iterative stimulation current pulses (supramaximal amplitude and duration indicated above) as follows: 2, 5, 15, 25, 40, 50, 75, 100 Hz. Each sequence of multiple pulses was applied, for EDL, for 1s followed by a relaxing period of 1s and, for Soleus, for 2s followed by a 1 s relaxing period. The absolute force was measured as amplitude at end of the pulse and normalized to the muscle weight (in N/g of muscle). The assessment of muscle fatigue was achieved by performing force-frequency relation protocol before (pre-fatigue) and 30 s after (post-fatigue) a fatigue protocol consisting of 30 successive sets of stimulations at 100 Hz.

### 2.6 Contraction measurement on isolated aortic rings

The thoracic aorta of rats was removed and placed into Krebs solution containing (in mM): 120 NaCl, 4.7 KCl, 2.5 CaCl_2_, 1.2 MgCl_2_, 1.2 KH_2_PO_4_, 15 NaHCO_3_, 11.1 D-glucose, pH 7.4. After separation of connective tissues, the thoracic segment of aorta was cut into rings of 3mm in length. Rat aorta rings were mounted between a fixed clamp at the base of a water-jacketed 5 ml organ bath contained an oxygenated (95% O_2_ and 5% CO_2_) Krebs solution and an IT1-25 isometric force transducer (Emka Technologies, Paris, France; (*13, 14*)). All experiments were performed at 37°C. A basal tension of 2 g was applied in all experiments. During 1 h, tissues were rinsed three times in Krebs solution and the basal tone was always monitored and adjusted to the range 400-1000 mg (*15*). 1 µM norepinephrine (denoted NE) was used to evoke the sustained contractile response.

### 2.7 Langendorff perfusion analyses

A left ventricular balloon system allows for real-time monitoring of the pressure developed by the contractile left ventricle of hearts mounted in a Langendorff set-up. To achieve this goal, control or HFHF rats were anaesthetized by intraperitoneal injection of sodium pentobarbital (60 mg/kg). The heart was quickly removed and the ascending aorta was connected according to the Langendorff technique (*16*) and a 0.06 ml latex balloon (VK 73-3479) was inserted in the left ventricle. The balloon was connected to a pressure transducer which was linked to the data acquisition system (PowerLab 425 - ADInstruments, Dunedin, New Zealand). Hemodynamic and functional parameters were recorded on a personal computer using LabChart software (ADInstruments, Dunedin, New Zealand). Hearts were allowed to stabilize during 30 minutes while perfused with standard Krebs solution. Functional parameters were assessed before, during and after a 30 minute-long ischemic insult. Preischemic period consisted in ventricular pressure monitoring in standard perfusion conditions during 10 minutes (Supplemental Fig. 14). Then, ischemia was induced by complete cessation of coronary flow during 30 minutes (global ischemia). At the end, reperfusion was initiated by re-establishing coronary flow and cardiac parameters were recorded and analyzed during the subsequent 30 minutes.

### 2.8 Statistical analysis

P values were calculated either by two-tailed *t-*tests or ANOVA, completed by adequate post-tests, as indicated the corresponding figure legends. All analyses were performed using the Graphpad Prism 5 software. ns: non-significant; ****P < 0.0001, ***P < 0.001, **P < 0.01 and *P < 0.05.

## 3. Results

### 3.1 Metabolic follow-up

After 5 weeks of diet, HFHF induced a significant increase in body weight (p < 0.05) maintained until the end of the study, in comparison to standard diet (CTL; Fig. 1A). Alterations in Glucose homeostasis were visible from the sixth week under HFHF (Fig. 1B). The HFHF rats also developed a dyslipidemia with a significant increase in blood of Triglyceride and Cholesterol levels as early as 2 and 4 weeks of age, respectively (Fig. 1C and 1D). These data recapitulated the observations from Lozano *et al.* (*9*). At 15 weeks, additional metabolic measurements were performed on 12 rats after prior gastric emptying (6 control and 6 HFHF; see below), and all animals were sacrificed to perform the phospholipid analyses and the *ex-vivo* experiments described below.

**Figure 1:**
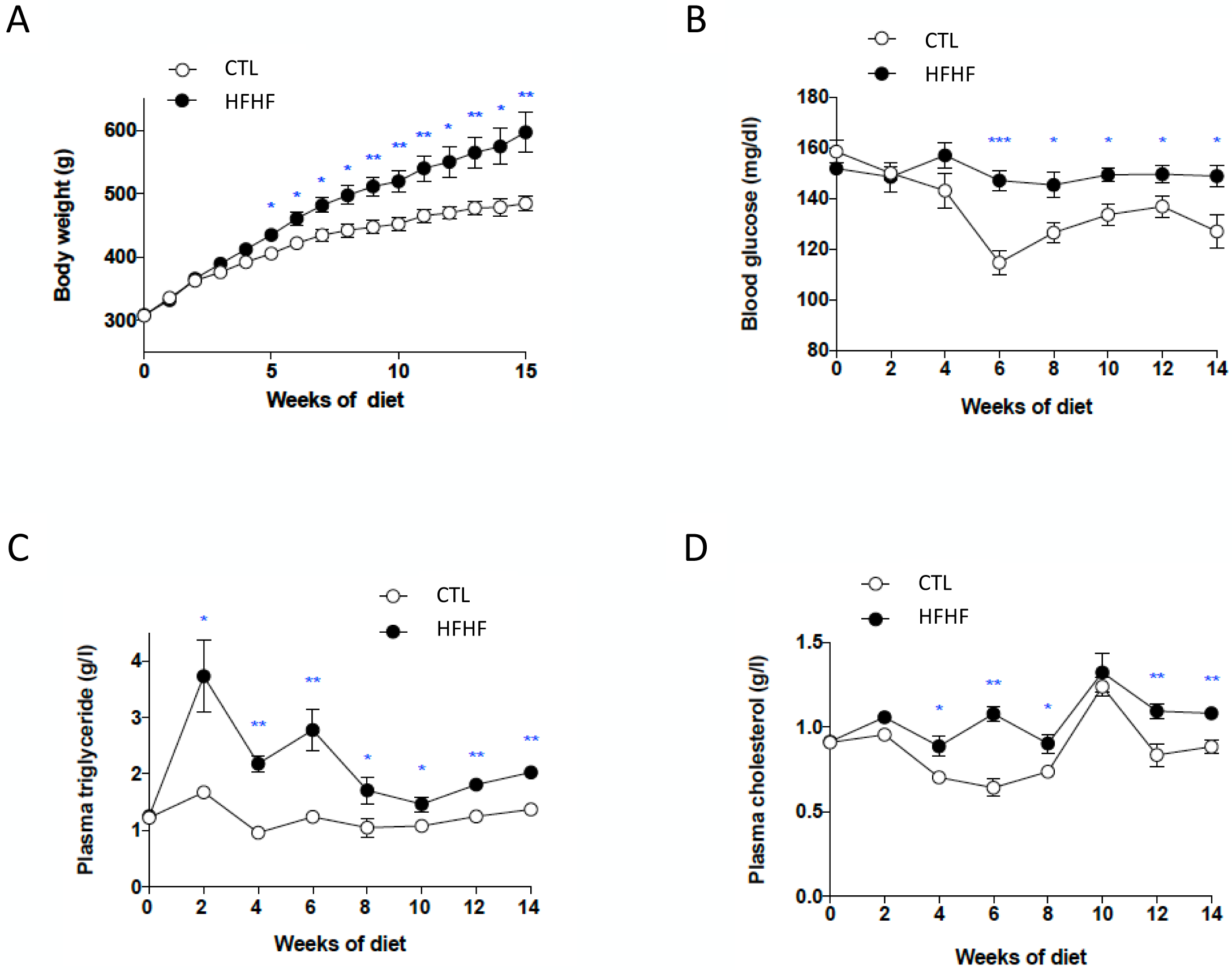
Longitudinal measurements (random fed conditions) performed during the study. During the longitudinal observation, body weight was measured each week (A) and plasma glucose (B), plasma Triglycerides (C) and plasma Cholesterol (D) levels were determined every 2 weeks at the same time of the day (2 PM) under random fed conditions, according to the protocols described in the “Materials and Methods” section, for CTL and HFHF rats (n = 8). Data are presented as means ± SD. Parameters were compared between CTL and HFHF rats using unpaired t-test.

### 3.2 Fatty acid distribution within phospholipids varies depending on the organ

In a first step, we determined the fatty acid distribution within the different phospholipid species (PL), namely PhosphatidylCholine (PC), PhosphatidylEthanolamine (PE), PhosphatidylSerine (PS) and PhosphatidylInositol (PI) in various organs from rats under a standard diet (CTL). This study was completed by the analysis of Plasmalogens which correspond to specific glycerophospholipids species containing a vinyl-ether bond at the *sn-*1 position (*4*). Among this lipid class, ethanolamine plasmalogens (PE(P)) are found in all rat organs ((*4*) and see below), whereas choline plasmalogens, which can be found in significant amounts in specific human tissues, are only detected as traces in rat organs ((*4*) and our unpublished data). The present study was performed on known targets of metabolic syndrome-induced disorders, such as the liver, skeletal muscles (EDL and Soleus), the vascular system (heart and aorta) and the pancreas, and was completed by the same analyses on the brain, spleen and lung. In this aim, total PL were extracted from the various organs and analyzed by mass spectrometry, in both the positive and negative ion modes, as described in the “Materials and Methods” section. Examples of characteristic spectra obtained in the negative and positive ion modes are displayed in Supplementary Fig. 1 and Fig. 2, respectively. The results obtained for PC are displayed in Fig. 2 as radar graphs and Supplementary Fig. 3 as histograms, and the data corresponding to PE, PS, PI and PE(P) are shown in Supplementary Fig. 4 to 7. In these figures, PL species are denominated by their initials followed by the total number of carbons and the number of carbon–carbon double bonds in their acyl chains (as an example, PC 38:4 corresponds to a PhosphatidylCholine bearing 38 carbon atoms and four double bonds in its acyl chains).

**Figure 2:**
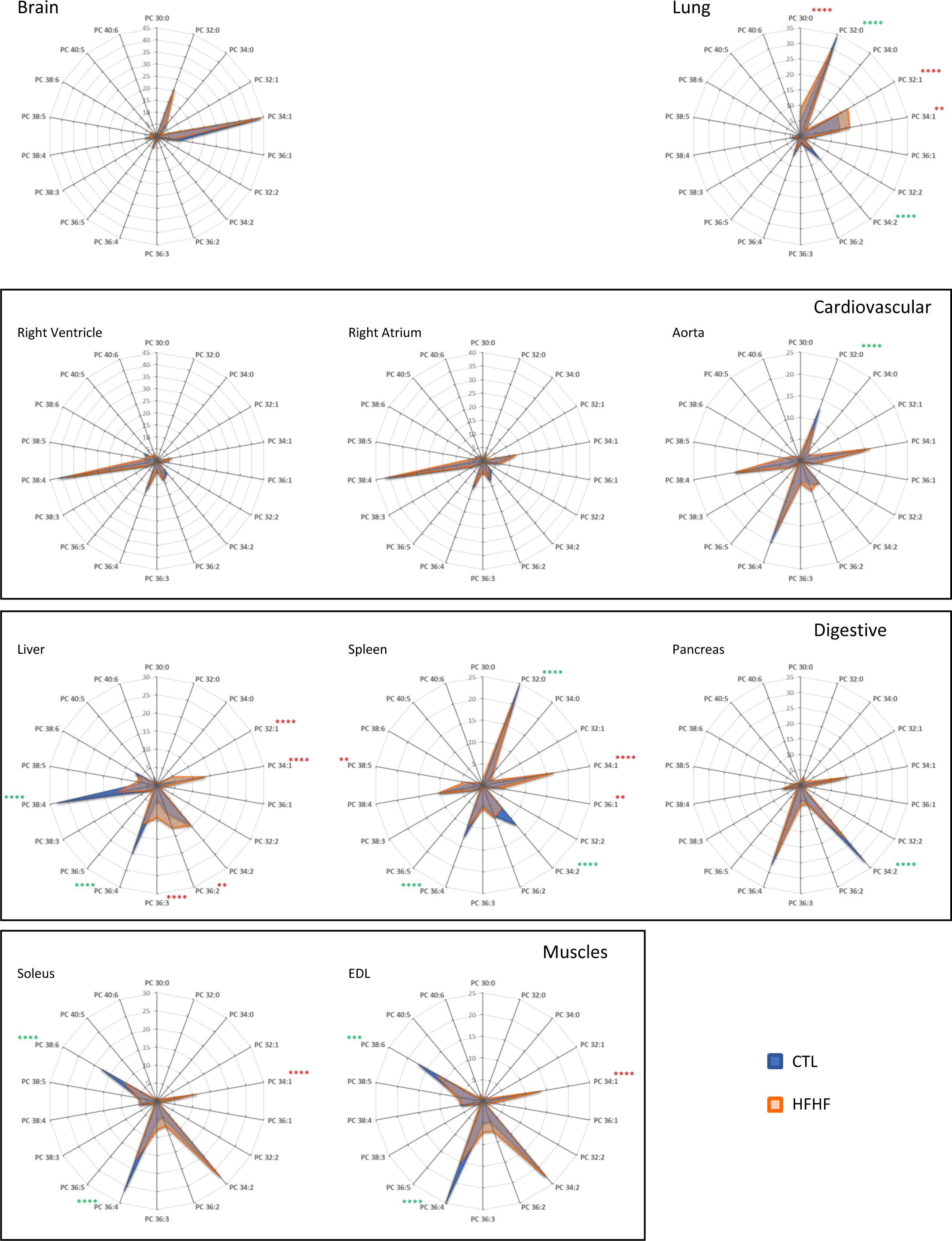
PC species distribution in various organs as a function of the diet. Total lipids were extracted and phospholipid species were purified and analyzed by ESI-MS from samples corresponding to the indicated organs obtained either rats fed with a normal (CTL) or HFHF diet (HFHF), as described in the “Materials and Methods”section. PC subspecies distribution in each case are displayed. The total carbon chain length (x) and number of carbon-carbon double bounds (y) of the main PC molecular species (x:y) are indicated. Values are means ± S.D. of four independent determinations from four individuals from both groups in each case. Statistical analysis was performed using two-way ANOVA, completed by Bonferroni post-tests to compare means variations between the two groups of animals for each PC subspecies. Significant differences between CTL and HFHF are indicated (****P < 0.0001, ***P < 0.001, and **P < 0.01), either in green if a specific subspecies is decreased under the HFHF diet as compared to CTL, or in red if this subspecies is increased under the HFHF regimen.

As already described in previous studies (*17*), a first observation is that PC is the PL species which displays the widest variations in terms of fatty acid chain distribution depending on the organ considered (Fig. 2 and Supplementary Fig. 3), whereas PI essentially appears as a major species (PI 38:4) in all the organs studied (Supplementary Fig. 4).

Concerning PC, it clearly appears that some organs are particularly enriched in species bearing polyunsaturated fatty acid chains (PUFA; more than two double-bonds/unsaturations in their fatty acid chains; *e. g.* PC 38:4), whereas others preferentially contain PC with two saturated fatty acyl chains (*e. g.* PC 32:0) (Fig. 2).

To visualize better these variations, the Double-Bond (DB) index was calculated in each case (Fig. 3A). As shown, the liver, skeletal muscles and the cardiovascular system were particularly enriched in PUFA-containing PC species (DB>2). By contrast, the spleen, the brain and the lung contained remarkably high amounts of saturated PC species (DB=0). The brain also differentiated from other organs by its high levels of monounsaturated PC species (DB=1). In the latter case, PC 34:1 appeared as the major species. Pancreas also displayed a very characteristic signature, with high levels of DB=2 PC.

**Figure 3:**
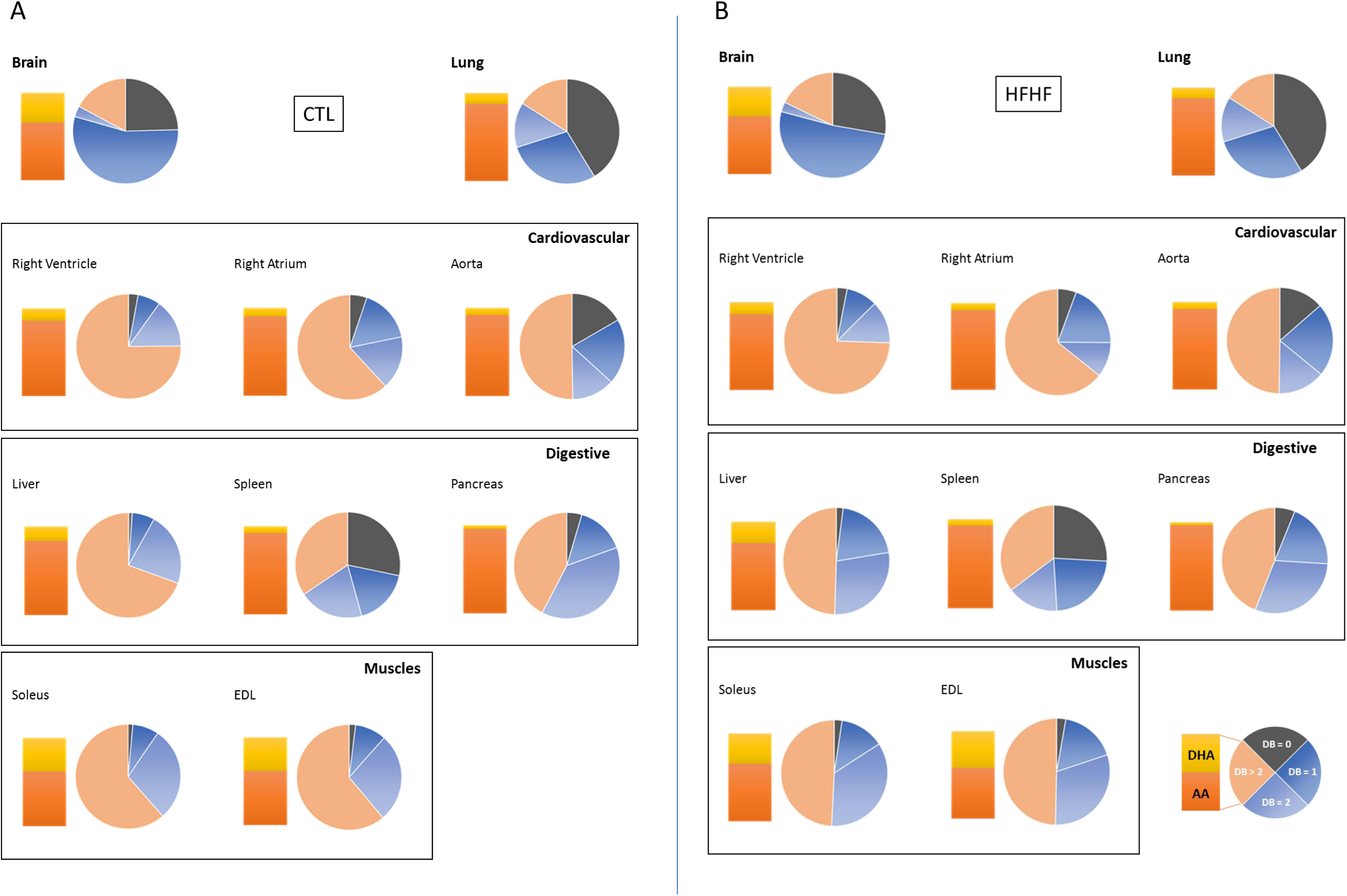
PC Double-Bond (DB) index and DHA to AA ratios in various organs as a function of the diet. Total lipids were extracted and phospholipid species were purified and analyzed by ESI-MS from samples corresponding to the indicated organs, obtained either rats fed with a normal (CTL) or HFHF diet (HFHF), as described in the “Materials and Methods” section. The relative percentage of saturated (DB=0: no double bonds) versus monounsaturated (DB=1: one double bond), diunsaturated (DB=2: two double bonds) and polyunsaturated (DB > 2: > two double bonds) phosphatidylcholine (PC) species was obtained from the PC subspecies distribution displayed in Fig. 2. The ratio of Docosahexaenoic Acid (DHA)- to Arachidonic Acid (AA)-containing PC subspecies in the various organs is also displayed.

Among PUFA, important variations could also be observed depending on the organ considered (Fig. 2 and Fig. 3A). If Arachidonic Acid (AA; 20:4) appeared as the most represented fatty acid in pancreas, spleen, liver, lung and the cardiovascular system, as combinations with Palmitate (PC 36:4) or Stearate (PC 38:4; Fig. 2; Table 1), Docosahexaenoic Acid (DHA; 22:6) was exquisitely enriched in skeletal muscles, in combination with Palmitate (PC 38:6; Fig. 2; Table 1).

**Table 1:** Distribution of the various PC subspecies in the liver and soleus as a function of the diet. PC studies were obtained after fragmentation studes (MS/MS) of the indicated PC species, as described in the “Materials and Methods” section and exemplified in Supplementary Fig. 2C. Subspecies that appeared to be increased or decreased under the HFHF diet as compared to the Normal diet (CTL) are indicated in Red and Green, respectively, based on the relative proportions of the various corresponding PC species obtained from direct infusion of lipid extracts, as displayed in Fig. 2. The relative quantities of the various subspecies corresponding to a given PC species are indicated by different font sizes.

|  | Liver |  |
| --- | --- | --- |
| PC species | CTL | HFHF |
| PC(32:1) | PC(16:0/16:1) | PC(16:0/16:1), PC(14:0/18:1) |
| PC(34:2) | PC(16:0/18:2) | PC(16:0/18:2), PC(16:1/18:1) |
| PC(34:1) | PC(16:0/18:1) | PC(16:0/18:1), PC(16:1/18:0) |
| PC(36:4) | PC(16:0/20:4) | PC(16:0/20:4), PC(18:2/18:2) |
| PC(36:3) | PC(16:0/20:3), PC(18:1/18:2) | PC(16:0/20:3), PC(18:1/18:2) |
| PC(36:2) | PC(18:0/18:2), PC(18:1/18:1) | PC(18:0/18:2), PC(18:1/18:1) |
| PC(38:4) | PC(18:0/20:4) | PC(18:0/20:4), PC(18:1/20:3) |
|  | Soleus |  |
|  | CTL | HFHF |
| PC(34:2) | PC(16:0/18:2) | PC(16:0/18:2) |
| PC(34:1) | PC(16:0/18:1) | PC(16:0/18:1) |
| PC(36:4) | PC(16:0/20:4), PC(18:2/18:2) | PC(16:0/20:4), PC(18:2/18:2) |
| PC(38:6) | PC(16:0/22:6), PC(18:2/20:4) | PC(16:0/22:6), PC(18:2/20:4) |

PE differentiated from PC in the sense that PUFA-containing species were systematically dominant, whatever the organ considered (Supplementary Fig. 5 and 9 A). However, the DHA to AA balance greatly varied among organs. As already observed for PC, DHA was particularly enriched in skeletal muscles, where PE 40:6 was the most represented species. Interestingly, DHA was also the most prominent PUFA in the brain, a situation very different of PC behavior in this organ (Fig. 2 and Fig. 3A). In the heart, whereas very similar signatures were observed in PC for ventricles and atria, PE 40:6 was enriched in the ventricles as compared to these latest. Even if less marked than with PC, the brain, spleen and lung appeared as the most saturated PE organs and the pancreas as the most DB=2 enriched organ.

The fatty acid distribution within PS paralleled the one observed with PE (Supplementary Fig. 6 and Supplementary Fig. 10 A), with PUFA being the major fatty acids in all the organs considered. Again, DHA was the most represented PUFA in skeletal muscles, brain, ventricle and atrium (PS 40:6), whereas AA was the most abundant in other organs (PS 38:4). The pancreas differentiated from others by a wider distribution of the fatty acyl chains, and relatively high levels of DB=2 species.

As already described elsewhere (*4*), PE(P) were detected in all the organs analyzed in this study, where they constituted between 30 and 56 % of ethanolamine-containing glycerophospholipids (*i. e.* PE and PE(P)), at the exception of the liver, in which their level dropped to 7 % only (data not-shown). Whatever the organ considered, PUFA-containing species were the most prominent (Supplementary Fig. 7 and 11A), with, as for PE and PS, a selective enrichment in DHA in the brain, the ventricle and skeletal muscles.

To summarize, all the organs studied here displayed very characteristic fatty acid distributions within PL and, therefore various saturation rates: they can be classified as DB=0/saturated organs (Spleen, lung), DB=1 (Brain), DB=2 (Pancreas) and DB>2 organs (liver, muscle and cardiovascular system). Among the latter, one can differentiate AA- (liver and Cardiovascular system) and DHA-enriched (skeletal muscle) organs. The brain is the organ which displays the most discrepancies between PL, PC being predominantly DB=2, whereas PE and PS are essentially DHA-enriched. These observations match and complete previous observations made by others (*17*).

### 3.3 Phospholipid species are differently affected by the High fat/High Fructose (HF/HF) diet, in an organ-specific manner

The high-fat diet used in this study is exquisitely enriched in SFA and MUFA, and specially Palmitate (16:0) and Oleate (18:1) (Table S1). Surprisingly, PE, PE(P), PS and PI all appeared to be quite insensitive to this oversupply, their overall fatty acid profile remaining similar whatever the diet (Supplementary Fig. 4 to 11).

By contrast, PC was the most affected class, but in a very selective organ-specific manner, the liver and the muscles being the most affected organs (Fig. 2 and 3B). In these tissues, DB=1 species appeared to be increased (PC 32:1 and PC 34:1 in the liver; PC 34:1 in muscles), at the expense of PUFA-containing species (PC 36:4 and PC 38:4 in the liver; PC 36:4 and PC 38:6 in muscles; Fig. 2). Notably, the same tendency was observed for PE and PS, but to lower levels and below significance (Supplementary Fig. 5, 6, 9 and 10).

Tandem MS experiments provided more information about fatty acid composition of PC species (for a representative example, see Supplementary Fig. 2C). As shown in Table 1, a closer look to MS/MS analyses revealed that, in muscles, PC species have the same composition of fatty acids (the same PC subspecies) in CTL and HFHF diets. Furthermore, MS results showed that PC(16:0/18:1) clearly accumulated at the expense of AA- (PC(16:0/20:4)) and DHA- (PC(16:0/22:6)) containing species under the HFHF diet (Fig. 2). This result reflected the fatty acid composition of the HFHF diet, which is not only enriched in 16:0 and 18:1, but also contains lower amounts of Linoleic (18:2) and Linolenic (18:3) acids, being respectively the precursors for AA (20:4) and DHA (22:6), than the standard diet.

By contrast, in the liver, the situation appeared to be more complex than expected. Indeed, increased amounts of PC 32:1 and PC 34:1 could be partly accounted in this organ by the apparition of PC species that were not detected in the liver of the rats under the standard diet, namely PC(14:0/18:1) and PC(16:1/18:0) (Table 1). Notably, 14:0, 16:1 and 18:0 are also enriched in the HFHF diet (Table S1). Morover, decreased amounts of PC 36:4 et PC 38:4 were not only related to a global decrease in AA-containing species, but were also accompanied with the formation of new species, namely PC(18:2/18:2) and PC(18:1/20:3), respectively (Table 1).

To summarize, two main organs appear to be highly reactive to the selective fatty acid-enrichment from the diet, namely the liver and skeletal muscles. Palmitate and Oleate preferentially distribute within PC under the form of PC 34:1. Moreover, a decrease in the amount of PUFA-containing species can also be observed in the same organs. This can be explained, at least for AA-containing species, by decreased amounts of the relevant precursor (namely Linoleic acid) in the HFHF diet.

### 3.4 Lipotoxicity in the liver

Among the various tissues, liver clearly appeared to be one of the main targets of diet-induced fatty-acid rearrangements in PL (Fig. 2 and 3). This was not a surprising observation, since the liver plays a key role in lipid metabolism, as the hub of fatty acid synthesis and lipid circulation through lipoprotein synthesis (*18*). In the HFHF model (*9*), alterations in liver function were manifested as early as 2 months following the shift to the enriched diet. The main manifestations at this early time point were the induction of steatosis, with a steatosis score of 1–2, according to Kleiner *et al.* (*19*), and increased hepatic levels of Reactive Oxygen Species (ROS). In the present study, we could also show that 4 months of HFHF resulted in a significant increase in liver weight (Fig. 4A), and in the amounts of circulating lipids, including Triglycerides (Fig. 4B), Cholesterol (Fig. 4C) and Non-Esterified Free Fatty Acids (NEFA; Fig. 4D). Analysis of the lipoprotein profile by Fast Protein Liquid Chromatography (FPLC) revealed that HFHF rats displayed increased plasma concentrations of lipoproteins rich in cholesterol (LDL and HDL; Fig. 4E). Moreover, we could also note an increase in Triglyceride levels within Chylomicrons / VLDL (fractions 3 to 10) and within LDL / HDLs (fractions 20 to 50, Fig. 4F).

**Figure 4:**
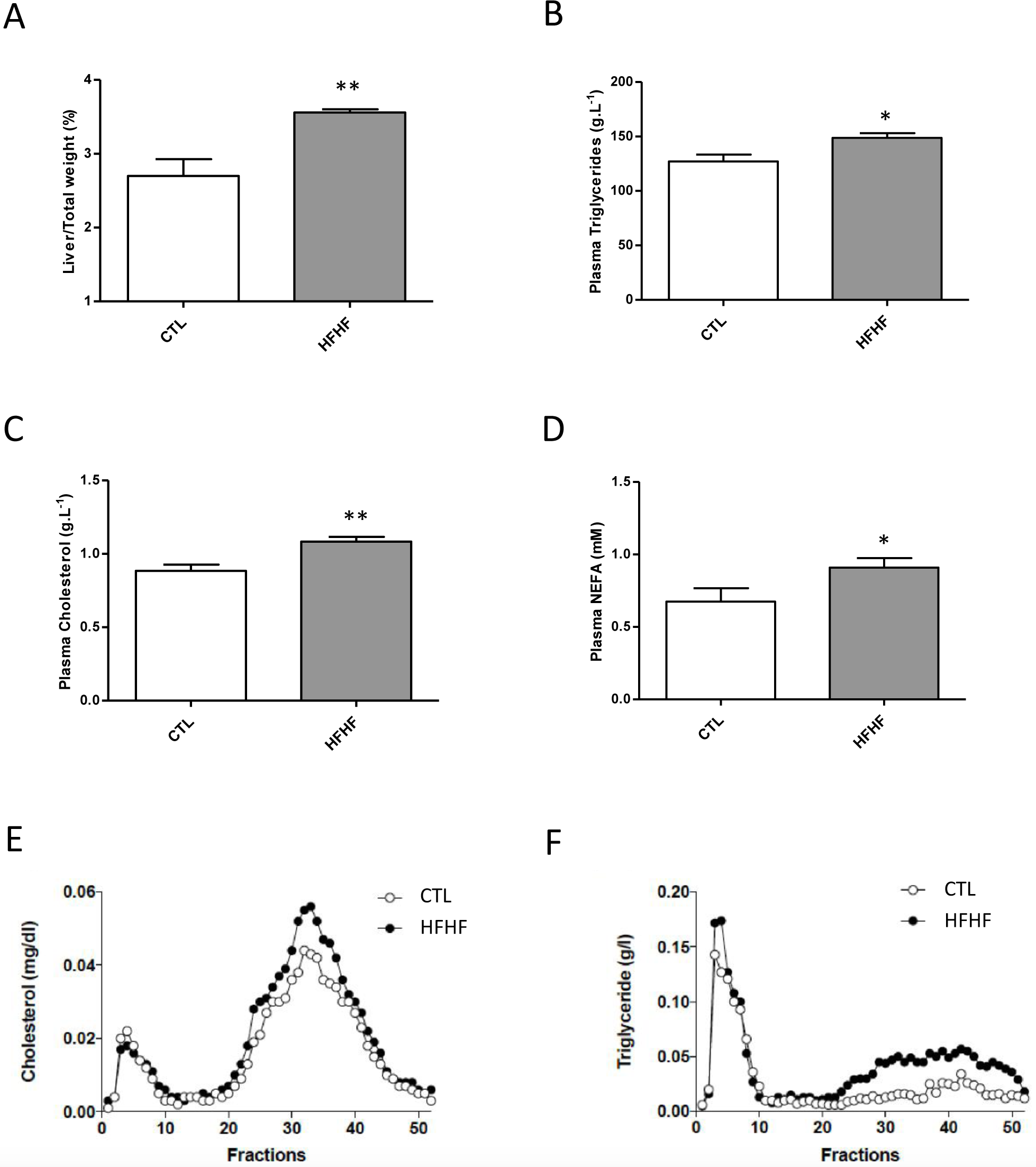
Impacts of the diet on liver function. Livers from rats fed with either a standard (CTL) or a HFHF diet (diet) for 15 weeks were dissected and weighted, and the Liver/Total weigth ratio was determined (A). 15 weeks after the initiation of the different diets, plasma Triglycerides (B), Cholesterol (C) and NEFA (D) levels were measured after a three hour fasting period to allow gastric emptying. In parallel, plasma samples were collected and subjected to fractionation by Fast Protein Liquid Chromatography (FPLC) and Cholesterol (E) and Triglyceride (F) concentrations in each fraction were measured. See the “Materials and Methods” section for details. All determinations were performed on 6 rats from each group. Values are means ± S.D. Parameters were compared between CTL and HFHF rats using unpaired *t*-test.

Since hepatic deposition of neutral lipids is a hallmark of steatosis, we also evaluated if such a deposition could be visualized in the HFHF model. In this aim, MS analyses were performed on non-purified lipid extracts from liver samples (Supplementary Fig. 12). As shown, clear-cut accumulations of Triglycerides (TG), Diglycerides (DG) and Free Cholesterol could be visualized in the liver of HFHF rats (Supplementary Fig. 12). These observations were confirmed with another analytical method, *i. e.* thin layer chromatography of hepatic neutral lipid fractions (Supplementary Fig. 13). These observations confirmed the previous data from Lozano et al. (*9*), showing a significant increase in TG in this organ as early as 2 months following induction of the HFHF diet, a situation which was maintained after 8 months. Interestingly, selective TG and DG species appeared to accumulate in the liver under HFHF, the main ones being TG(52:2), TG(54:5), DG(34:1) a,d DG(36:2) (Supplementary Fig. 12). Complementary MS/MS analyses (our unpublished data) revealed that these lipids corresponded to 16:0- and 18:1-containing species, namely TG(16:0/18:1/18:1), TG(16:0/18:1/20:4), DG(16:0/18:1) and DG(18:1/18:1), therefore reflecting the fatty acid composition of the HFHF diet.

### 3.5 Lipotoxicity and muscle function

With the liver, muscles were the most affected tissues in terms of their sensitivity to fatty acid rearrangements within PL (Fig. 2 and 3). Interestingly however, by contrast to liver, this redistribution was not paralleled by the deposition of neutral lipids (TG, DG and Cholesterol; Supplementary Fig. 12 and 13). Fatty acid rearrangements corresponded to a decrease in the amounts of DHA containing PC species (namely PC 38:6), a lipid species that was exquisitely enriched in these tissues (Fig. 2; Table 1). Notably, the same observations could be done in either the fast-twitch (EDL) and slow-twitch (Soleus) types of muscles (Fig. 2).

Obesity can cause a decline in contractile function of skeletal muscle. Isolated muscle preparations show that obesity often leads to a decrease in force produced per muscle cross-sectional area, and power produced per muscle mass (*20*).

We therefore investigated further the effects of HFHF diet on the functional features of both fast-twitch and slow twitch skeletal muscles. In this aim, the fast-twitch extensor digitorum longus muscles (EDL) and the slow twitch Soleus muscles (Soleus) were isolated from rats under both diets. A first observation was that absolute mass of these skeletal muscles was increased under the HFHF diet (Fig 5A and 5B).

**Figure 5:**
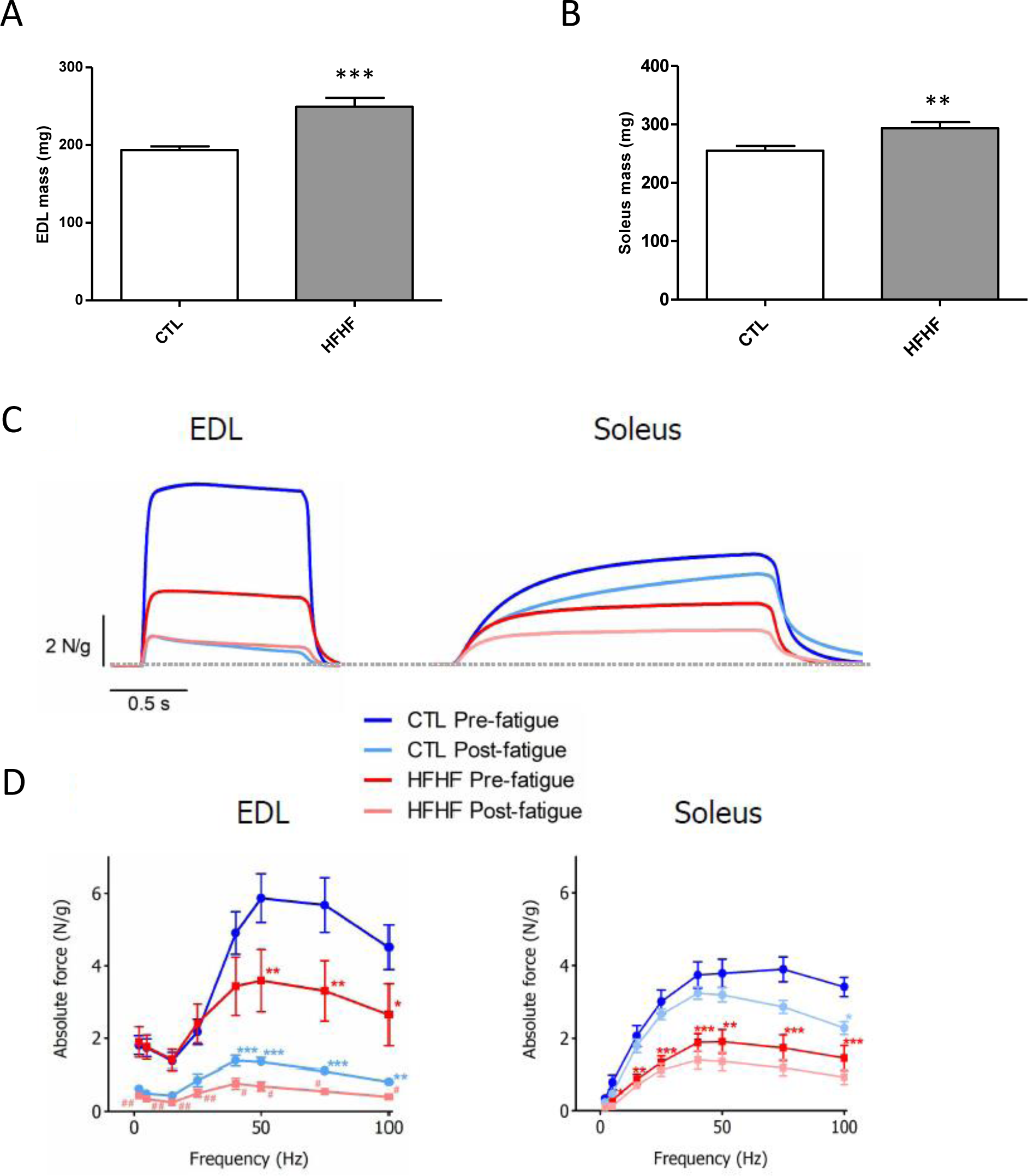
Impacts of the diet on the function of skeletal muscles. 15 weeks after the initiation of the different diets (CTL or HFHF), plasma Glucose (A) and Insulin (B) levels were measured after a three hour fasting period to allow gastric emptying, as described under the “Materials and Methods” section (n = 6). At the same time point, the EDL (C) and Soleus (D) muscles were dissected and their weigth was determined for comparison between CTL and HFHF rats (n = 11). Values are means ± S.E.M. Parameters were compared between CTL and HFHF rats using unpaired *t*-test (A-D). Effects of HFHF diet on tetanus amplitude and fatigue of EDL and Soleus muscle (F and G). Examples of tetanus responses to electrical field stimulation at 100 Hz for EDL (left panel) and Soleus (right panel) before and after a fatigue protocol in control (blue traces) and HFHF (red traces) rats (F). Force-frequency relationships for the same types of muscle than in F (G). Values are means ± S.E.M. Statistical tests were performed using one-way analysis of variance and a Dunnett’s multiple comparison as post test.

In a next step, Soleus and EDL were stimulated with field electrodes to measure force characteristics in two different states: before fatigue (pre-fatigue) and immediately after a fatigue protocol (post-fatigue) (Fig. 5 C-D).

Figure 5C displays examples of tetanus force responses, relative to muscle mass, in EDL and Soleus muscles from control CTL and HFHF rats in the two pre- and post-fatigue stimulation conditions. In these examples, force was classically found weaker in CTL Soleus muscle than in EDL CTL muscles in pre-fatigue conditions (mean values: EDL: 4.5 ± 0.6, Soleus 3.4 ± 0.3 N/g). Tetanic force was found impaired by HFHF diet in EDL muscles and a reduction, but with less impact, was also observed in Soleus (Fig. 5C). When we analyzed the force-frequency curves (Fig. 5D), for both EDL and Soleus muscles, the HFHF diet (red curves) resulted in a decreased muscle tetanic force, compared with CTL, starting at stimulation frequencies greater than 40 Hz - 50 Hz (At 100 Hz, EDL: 2.6 ± 0.8 N/g in HFHF versus 4.5 ± 0.6 N/g in the CTL group, Soleus: 1.5 ± 0.3 N/g in HFHF versus 3.4 ± 0.3 N/g in the CTL group). These findings indicate that a HFHF diet impairs contractile force in both fast-twitch and slow twitch muscles.

Moreover, a significant impairment of tetanic force after fatigue protocols was observed. At 100 Hz, in CTL EDL muscle, fatigue protocol led to a 82 % decrease (from 4.5 ± 0.6 to 0.8 ± 0.1 N/g) of tetanic force and such a decrease reached 85 % in HFHF EDL (from 2.6 ± 0.8 to 0.4 ± 0.1 N/g). The same effect, at a lesser extent, was recorded in Soleus: 32 % (from 3.4 ± 0.3 to 2.3 ± 0.2 N/g) in CTL and 39% (from 1.5 ± 0.3 to 0.9 ± 0.2 N/g) in HFHF.

To conclude, adaptations occurring in response to HFHF diet result in a general muscle force loss consistent with that observed in humans (*21*). These findings would imply that HFHF diet induces a drastic decrease of the tetanus force whatever the type of muscle and without significantly changing the behaviour towards fatigue.

### 3.5 Lipotoxicity and the cardiovascular system

By contrast to the liver and skeletal muscles, the cardiovascular system appeared to be quite protected from diet-induced fatty acid redistribution within PL (Fig. 2 and Fig. 3).

However, since many evidences exist on the relationship between obesity and cardiovascular disease (CVD) in humans, even if the degree and the duration of obesity appears to affect the severity of CVD (*22*), we decided to evaluate further the impacts of the HFHF diet on the cardiovascular system.

First, a maximal exercise test was used as an indicator of the cardio-respiratory capacity of rats. Interestingly, the Maximum Running Speeds (MRS) of the CTRL and the HFHF group were not significantly different post-diet, with values of 27.8 ± 0.83 and 29.4 ± 0.61 m/min, respectively (Fig. 6A). Therefore, HFHF clearly did not impair the global functional capacity of the animals.

**Figure 6:**
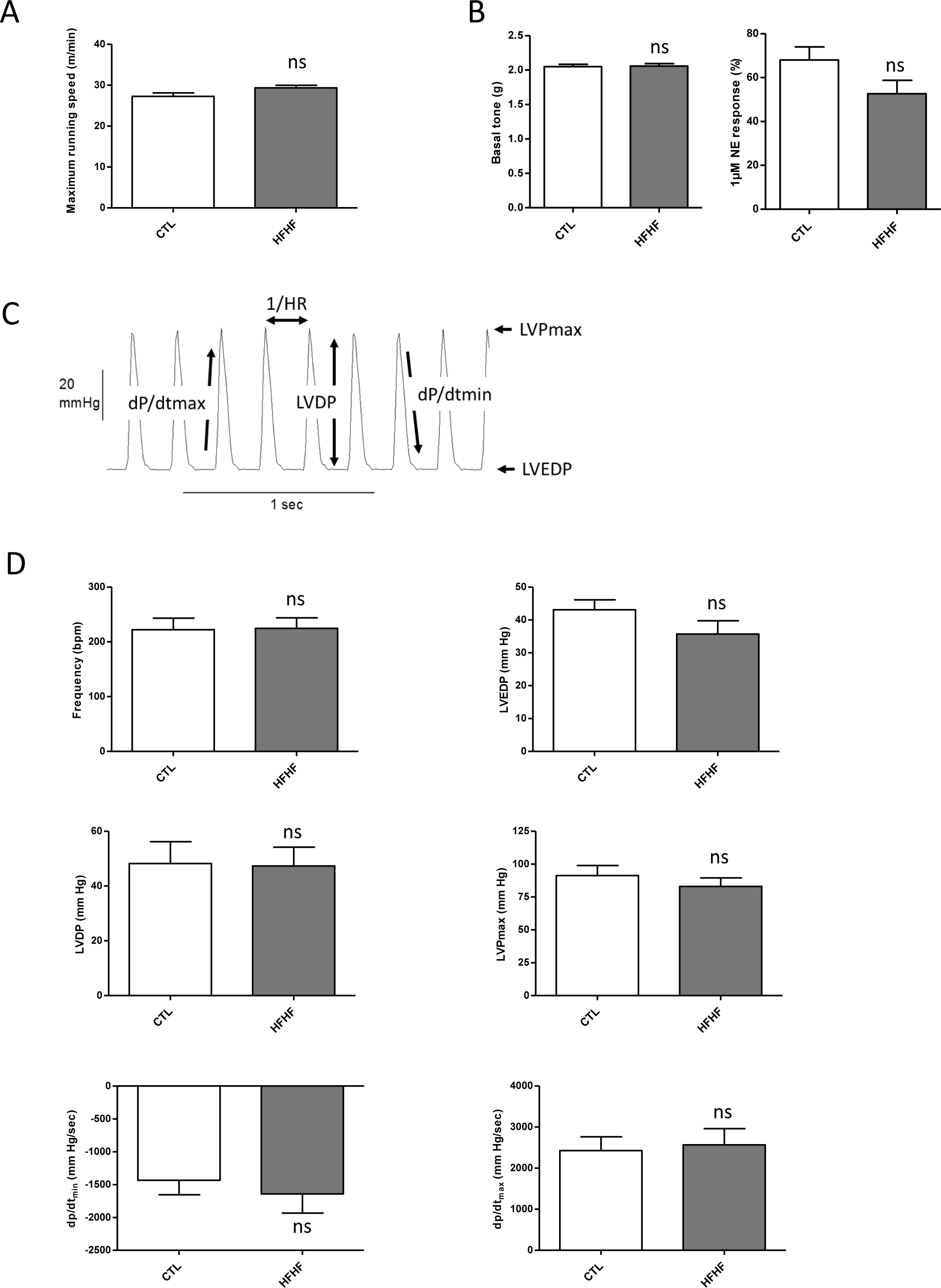
Impacts of the diet on the Cardiovascular System. The Maximal Running Speed (MRS) was determined 15 weeks after the initiation of the different diets (CTL (n = 7) or HFHF (n = 8)) (A). At the same time point, the basal tone and the induced-contraction was measured on rat aorta rings (B). Aorta rings obtained from four control and five HF/HF rats were mounted between a fixed clamp and incubated in Krebs solution to determine the basal tone (Left panel). 1 µM Norepinephrine (denoted NE) was added to the same aorta rings to evoke the sustained contractile response (Right panel). The pressure developed by the contractile left ventricle of the animals was also determined using a Langendorff set-up (C and D). Rat hearts from either control (n=11) or HFHF (n=12) groups were submitted to the protocol illustrated in Supplementary Fig. 10. Parameters recorded during the whole protocol are illustrated in C). The results obtained during pre-ischemic period are presented in D) as mean ± SEM. Pre-ischemic parameters were compared between CTL and HFHF rats using unpaired *t*-test.

Obese subjects with insulin resistance and hypertension have abnormal aortic elastic function, which may predispose them to the development of left ventricular dysfunction (*23*). In this context, we compared the basal tone on control and HFHF rat aorta, but we observed no difference between the two types of aorta rings (Fig. 6B). The norepinephrine (denoted NE)-induced vasoconstriction was also similar on control and HF/HF rat rings (Fig. 6B).

Direct cardiac structural abnormalities and alterations in ventricular function have been shown to occur in severely obese patients and in a process that may predispose them to heart failure. More specifically, left ventricular (LV) hypertrophy in severe obesity (either eccentric or concentric) is frequently observed and the direct implication of metabolic disturbance, including Lipotoxicity, has been suggested (*22*).

The left ventricular balloon system allows for real-time monitoring of the pressure developed by the contractile left ventricle of hearts mounted in a Langendorff set-up. Rat hearts from either control (n=11) or HFHF (n=12) groups were submitted to the protocol illustrated in Supplementary Fig. 14. Parameters recorded during the pre-ischemic period (Fig. 6C) are illustrated in Fig. 6D, which shows that no significant difference was found between the two groups, whatever the parameter under consideration. In other words, the contractile behavior of rat hearts was not impacted by the feeding diet imposed to rats during the 15 weeks preceding the experiment.

When the hearts from both groups were submitted to global ischemia (Supplementary Fig. 15), contractile performances rapidly decreased until developed pressure completely vanished. Again, no significant difference was observed between control and HFHF rats, whatever the recorded parameter. When perfusion was restored, as illustrated in Supplementary Fig. 15, cardiac performance was rapidly recovered. It seems that there is a tendency of hearts from control rats to recover better than HFHF hearts during reperfusion. However, this tendency only reaches statistical significance for LVDP at 1 and 5 min following reperfusion onset, all the other values remaining non-significantly different between the two groups (Supplementary Fig. 15).

To conclude, the cardiovascular system appeared to be functionally protected during the early onset of the metabolic syndrome under the HFHF diet in our conditions.

## 4. Discussion

It is known since a long time that the fatty acid composition of the diet can influence Phospholipid signature in various tissues (For review, see (*24*)). However, an important conclusion from the present study is that not all organs are equal to this redistribution. Indeed, in the HFHF rat model, most organs were protected from fatty acid rearrangements, at least during the early onset of the metabolic syndrome, with the liver and skeletal muscles being the preferred targets (Fig. 2 and Fig. 3). A second important conclusion of the present study is that fatty acids from the diet can distribute within PL in a very selective way: PC appears to be the preferred sink for this distribution.

In the liver, this distribution was paralleled by the deposition of neutral lipids, including Di- and Tri-glycerides, Cholesterol and Steryl-esters (Supplementary Fig. 12 and 13). Hepatic deposition of neutral lipid is a hallmark of dyslipidemia in obesity and it has been proposed to promote hepatic insulin resistance associated with nonalcoholic fatty liver disease (NAFLD), which is a major factor in the pathogenesis of type 2 diabetes and the metabolic syndrome (*25*, 26). As observed for PC, the main TG and DG species which accumulated in this organ corresponded to the ones containing the fatty acids which are specifically enriched in the HFHF diet used in this study, *i. e.* 16:0 and 18:1 (Supplementary Table 1): PC(16:0/18:1) (Table 1), TG(16:0/18:1/18:1), TG(16:0/18:1/20:4), DG(16:0/18:1) and DG(18:1/18:1) (Supplementary Fig. 12). Interestingly, in obese individuals, DG can inhibit insulin signaling by activation of Protein Kinase C (PKC) isoforms (*26*). In these patients, and as in the HFHF model, hepatic DG composed of 16:0/18:1 and 18:1/18:1 are most abundant and also strongly related with insulin resistance (*26*). Accordingly, HFHF resulted in the present study in a significant increase in the concentrations of circulating Glucose (Fig. 1B) and Insulin (data not-shown), confirming previous data from Lozano *et al*. (*9*) showing that after 2 months, the HOMA2-IR (homeostasis model assessment) values were higher than 2.4 in HFHF rats. These data demonstrated insulin resistance in this model, at a very early stage in the onset of the metabolic syndrome. To summarize, similar lipid depositions/rearrangements are observed in the liver of obese individuals and of HFHF rats, with likely similar impacts on the initiation of the insulin-resistance phenotype.

If the impacts of HFHF on the liver were not a a surprising observation, since this organ plays a key role in lipid metabolism, the fact that skeletal muscles were the second most affected tissues in terms of PC fatty acid rearrangements was less predictable (Fig. 2). Importantly, by contrast to liver, this redistribution was not paralleled by deposition of neutral lipids in these tissues (Supplementary Fig. 12 and 13).

Obesity is generally associated with changes in muscle quality, as it appears to result in larger muscles of lower quality (*i. e.* less contractile force per unit of cross sectional area and lower power output per unit of muscle mass), which have the same absolute force and power output of smaller muscles in lean individuals (*20*). The exact same observations were made in the present study, HFHF resulting in absolute increases in EDL and Soleus masses (Fig. 5A and 5B), but reduced force per muscle mass (Fig. 5C and 5D). Increase in muscle mass likely compensates poor muscle quality, at least in the early onset of the metabolic syndrome, as manifested by the similar performances of the HFHF rats in the maximal exercise tests (Fig. 6A). If the connection between muscle force and the observed decrease in PUFA-containing PL remains correlative at this step, knowing the importance of such lipid species in membrane plasticity/elasticity (*6*, 27), additional experiments aiming at studying the intimate relationships between the levels of PUFA-enriched PL, the membrane properties of muscle cells and their ability to stretch/contract, will undoutfully shed new lights on these mechanisms.

Finally, the fact that the cardiovascular system (CS) was protected from fatty-acid rearrangements within PL and remained largely unaffected on a functional point of view was also an unexpected result. These observations suggest that protective mechanisms under dyslipidemia do exist to channel excess fatty acids to skeletal muscles rather than to CS. It has been demonstrated that the prognosis of Cardiovascular diseases (CVD) of a patient who just became obese might be different from another who has been obese for many yeasr (*22*). In a pionnering study, Nakajima *et al.* (*28*) demonstrated that alterations of cardiac performance in obese individuals is attributed not only to the excess of body weight but also to the duration of obesity. Long lasting experiences to evaluate the impacts of HFHF on the CS, both on PL fatty acid signature and overall performance will undoutfully help to establish clearer connections between these processes.

To conclude, this study is, to our knowledge, the first of its kind to give such an overview of the distribution of fatty acids originating from the diet within PL in various organs and their functional performances. Further studies aiming at establishing direct links between PL fatty acid composition, relevant membrane properties in targeted cells, and organ function, particularly in the later steps of the metabolic syndrome, will undoutfully help at understanding the impacts of lipotoxicity on the progression of the associated diseases/comorbidities in this complex pathology.

## Acknowledgements

This work has benefited from the facilities and expertise of the Therassay facility (University of Nantes, France). In this context, Dr. Cédric Le May and Dr. Maud Chétiveaux are acknowledged for their precious help and expertise in the determination of the metabolic parameters displayed in this study. The team from the PreBios facility (University of Poitiers, France) is also acknowledged for taking care of the animals during the course of this study. This work was supported by the Ministère de l’Education Nationale, de l’Enseignement Supérieur et de la Recherche (French MENRT) and the FEDER (Fonds Européen de Développement Régional), with grants to L. K. and A. B.

## References

1. Kusminski, C., Shetty, S., Orci, L., Unger, R., and Scherer, P. (2009) Diabetes and apoptosis: lipotoxicity, Apoptosis 14, 1484–1495.

2. Cascio, G., Schiera, G., and Liegro, I. D. (2012) Dietary Fatty Acids in Metabolic Syndrome, Diabetes and Cardiovascular Diseases, Current Diabetes Reviews 8, 2–17.

3. Christie, W. W. (1985) Rapid separation and quantification of lipid classes by high performance liquid chromatography and mass (light-scattering) detection, Journal of Lipid Research 26, 507–512.

4. Braverman, N. E., and Moser, A. B. (2012) Functions of plasmalogen lipids in health and disease, Biochimica et Biophysica Acta (BBA) - Molecular Basis of Disease 1822, 1442–1452.

5. Honsho, M., and Fujiki, Y. (2017) Plasmalogen homeostasis – regulation of plasmalogen biosynthesis and its physiological consequence in mammals, FEBS Letters 591, 2720–2729.

6. Antonny, B., Vanni, S., Shindou, H., and Ferreira, T. (2015) From zero to six double bonds: phospholipid unsaturation and organelle function, Trends in Cell Biology 25, 427–436.

7. Bacle, A., and Ferreira, T. (2019) Strategies to Counter Saturated Fatty Acid (SFA)-Mediated Lipointoxication, In The Molecular Nutrition of Fats (Patel, V. B., Ed.), pp 347–363, Academic Press.

8. Wong, S. K., Chin, K.-Y., Suhaimi, F. H., Fairus, A., and Ima-Nirwana, S. (2016) Animal models of metabolic syndrome: a review, Nutrition & Metabolism 13, 65–65.

9. Lozano, I., Van der Werf, R., Bietiger, W., Seyfritz, E., Peronet, C., Pinget, M., Jeandidier, N., Maillard, E., Marchioni, E., Sigrist, S., and Dal, S. (2016) High-fructose and high-fat diet-induced disorders in rats: impact on diabetes risk, hepatic and vascular complications, Nutrition & Metabolism 13, 15–15.

10. Kadri, L., Ferru-Clément, R., Bacle, A., Payet, L.-A., Cantereau, A., Hélye, R., Becq, F., Jayle, C., Vandebrouck, C., and Ferreira, T. (2018) Modulation of cellular membrane properties as a potential therapeutic strategy to counter lipointoxication in obstructive pulmonary diseases, Biochimica et Biophysica Acta (BBA) - Molecular Basis of Disease 1864, 3069–3084.

11. Husen, P., Tarasov, K., Katafiasz, M., Sokol, E., Vogt, J., Baumgart, J., Nitsch, R., Ekroos, K., and Ejsing, C. S. (2013) Analysis of Lipid Experiments (ALEX): A Software Framework for Analysis of High-Resolution Shotgun Lipidomics Data, PLoS ONE 8, e79736.

12. Carneiro-Júnior, M. A., Quintão-Júnior, J. F., Drummond, L. R., Lavorato, V. N., Drummond, F. R., da Cunha, D. N. Q., Amadeu, M. A., Felix, L. B., de Oliveira, E. M., Cruz, J. S., Prímola-Gomes, T. N., Mill, J. G., and Natali, A. J. (2013) The benefits of endurance training in cardiomyocyte function in hypertensive rats are reversed within four weeks of detraining, Journal of Molecular and Cellular Cardiology 57, 119–128.

13. Fortner, C. N., Lorenz, J. N., and Paul, R. J. (2001) Chloride channel function is linked to epithelium-dependent airway relaxation, Am J Physiol Lung Cell Mol Physiol 280, 334–341.

14. Vandebrouck, C., Melin, P., Norez, C., Robert, R., Guibert, C., Mettey, Y., and Becq, F. (2006) Evidence that CFTR is expressed in rat tracheal smooth muscle cells and contributes to bronchodilation, Respiratory Research 7, 113.

15. Watson, N., Magnussen, H., and Rabe, K. F. (1998) The relevance of resting tension to responsiveness and inherent tone of human bronchial smooth muscle, British Journal of Pharmacology 123, 694–700.

16. Bell, R. M., Mocanu, M. M., and Yellon, D. M. (2011) Retrograde heart perfusion: The Langendorff technique of isolated heart perfusion, Journal of Molecular and Cellular Cardiology 50, 940–950.

17. Hicks, A. M., DeLong, C. J., Thomas, M. J., Samuel, M., and Cui, Z. (2006) Unique molecular signatures of glycerophospholipid species in different rat tissues analyzed by tandem mass spectrometry, Biochimica et Biophysica Acta (BBA) - Molecular and Cell Biology of Lipids 1761, 1022–1029.

18. Nguyen, P., Leray, V., Diez, M., Serisier, S., Bloc’h, J. L., Siliart, B., and Dumon, H. (2008) Liver lipid metabolism, Journal of Animal Physiology and Animal Nutrition 92, 272–283.

19. Kleiner, D. E., Brunt, E. M., Van Natta, M., Behling, C., Contos, M. J., Cummings, O. W., Ferrell, L. D., Liu, Y.-C., Torbenson, M. S., Unalp-Arida, A., Yeh, M., McCullough, A. J., and Sanyal, A. J. (2005) Design and validation of a histological scoring system for nonalcoholic fatty liver disease, Hepatology 41, 1313–1321.

20. Tallis, J., James, R. S., and Seebacher, F. (2018) The effects of obesity on skeletal muscle contractile function, The Journal of Experimental Biology 221.

21. Sayer, A. A., Dennison, E. M., Syddall, H. E., Gilbody, H. J., Phillips, D. I. W., and Cooper, C. (2005) Type 2 Diabetes, Muscle Strength, and Impaired Physical Function, The tip of the iceberg? 28, 2541–2542.

22. Ortega Francisco, B., Lavie Carl, J., and Blair Steven, N. (2016) Obesity and Cardiovascular Disease, Circulation Research 118, 1752–1770.

23. Robinson, M. R., Scheuermann-Freestone, M., Leeson, P., Channon, K. M., Clarke, K., Neubauer, S., and Wiesmann, F. (2008) Uncomplicated obesity is associated with abnormal aortic function assessed by cardiovascular magnetic resonance, Journal of cardiovascular magnetic resonance: official journal of the Society for Cardiovascular Magnetic Resonance 10, 10–10.

24. Gimenez, M. S., Oliveros, L. B., and Gomez, N. N. (2011) Nutritional deficiencies and phospholipid metabolism, International journal of molecular sciences 12, 2408–2433.

25. Klop, B., Elte, J. W. F., and Cabezas, M. C. (2013) Dyslipidemia in obesity: mechanisms and potential targets, Nutrients 5, 1218–1240.

26. Kumashiro, N., Erion, D. M., Zhang, D., Kahn, M., Beddow, S. A., Chu, X., Still, C. D., Gerhard, G. S., Han, X., Dziura, J., Petersen, K. F., Samuel, V. T., and Shulman, G. I. (2011) Cellular mechanism of insulin resistance in nonalcoholic fatty liver disease, Proceedings of the National Academy of Sciences 108, 16381–16385.

27. Pinot, M., Vanni, S., Pagnotta, S., Lacas-Gervais, S., Payet, L.-A., Ferreira, T., Gautier, R., Goud, B., Antonny, B., and Barelli, H. (2014) Polyunsaturated phospholipids facilitate membrane deformation and fission by endocytic proteins, Science 345, 693–697.

28. Nakajima, T., Fujioka, S., Tokunaga, K., Hirobe, K., Matsuzawa, Y., and Tarui, S. (1985) Noninvasive study of left ventricular performance in obese patients: influence of duration of obesity, Circulation 71, 481–486.

