## Supplementary Text for "A comprehensive study of Phospholipid fatty acid rearrangements in the early onset of the metabolic syndrome: correlations to organ dysfunction"

### Supplementary Data

#### Supplementary Figure 1: Analysis strategy of Phospholipids (PI, PE and PS) by mass spectrometry in the negative ion mode.

A) Mass spectrum of liver lipid extract in negative ion mode showing main phospholipid species as $[M-H]^-$ ion. B) MS/MS spectrum of PI (38:4) in negative ion mode showing different fragment ions. Characteristic ions with m/z 259.2, 241 and 223 corresponding to the inositol phosphate ion, inositol phosphate ion with the loss of one and two molecules of water respectively, allowed the identification of PI polar head group. Abundant ions with m/z 283.2 (stearic acid detected as $RCOO^-$) and m/z 303.2 (arachidonic acid detected as $RCOO^-$) allowed the identification of the fatty acyl chains. Other fragment ions with m/z: 78.9 ($PO_3^-$ ion), 153 (glycerol-3-phosphate ion with loss of $H_2O$), 419.2 (neutral loss of sn2 RCOOH group and inositol from $[M-H]^-$), 439.2 (neutral loss of sn1 RCOOH group and inositol from $[M-H]^-$), 581.3 (neutral loss of sn2 RCOOH group from $[M-H]^-$) and 599.3 (loss of sn2 acyl chain as (RCH=C=O) from $[M-H]^-$) were also detected in negative ion mode. C) MS/MS spectrum of PE (38:4) in negative ion mode. Characteristic ions of m/z 140 (ethanolamine phosphate ion) and m/z 196 (dilyso-PE with the loss of $H_2O$) allowed the identification of PE polar head group. Abundant ions of m/z 283.2 (stearic acid detected as $RCOO^-$) and m/z 303.2 (arachidonic acid detected as $RCOO^-$) allowed the identification of the fatty acyl chains. Other ions corresponding to the neutral loss of sn2 as RCOOH group from $[M-H]^-$ and the loss of sn2 acyl chain as RCH=C=O from $[M-H]^-$, with m/z 462.3 and 480.3 respectively, were also detected. D) MS/MS spectrum of PS (38:4) in negative ion mode. Characteristic fragment ion corresponding to the loss of serine from precursor ion with m/z 723.5 was detected. Fragment ions corresponding to the loss of sn1 acyl chain as RCH=C=O and serine from $[M-H]^-$, neutral loss of sn1 RCOOH group and serine from $[M-H]^-$, and neural loss of sn2 RCOOH group and serine from $[M-H]^-$ with m/z 457.2, 439.2 and 419.2 respectively, were observed to be characteristic fragments of this compound in negative ion mode. Fragment ions with m/z 283.2 (stearic acid detected as RCOO-) and m/z 303.2 (arachidonic acid detected as RCOO-) allowed the identification of the fatty acyl chains. Glycerol-3-phosphate ion with loss of $H_2O$ (m/z 152.9) was also detected in negative ion mode. Data processing of fragment ions spectra was carried out with Biovia Draw 19,1© and MassLynx 4,1© software with the help of Lipid Maps Lipidomics Gatway®.

#### Supplementary Figure 2: Analysis strategy of Phosphatidylcholine species by mass spectrometry.

A) Mass spectrum of liver lipid extract in positive ion mode showing main PhosphatidylCholine (PC) species as protonated ions $[M+H]^+$. B) MS/MS spectrum of PC (38:4) in positive ion mode. This spectrum allowed the identification of the polar head of PC class (characteristic fragment ion with m/z 184). C) MS/MS spectrum of PC (38:4) in negative ion mode as $[M+Cl]^-$ adduct. Abundant ions with m/z 283.2 (stearic acid detected as $RCOO^-$) and m/z 303.2 (arachidonic acid detected as $RCOO^-$) allowed the identification of the fatty acyl chains. Other characteristic fragment ions corresponding to the loss of $CH_3$ and chloride from precursor ion, and to the loss of sn2 acyl chain as RCH=C=O, $CH_3$ and chloride from precursor ion with m/z 794.5 and 508.3 respectively, were detected in negative ion mode. Ion corresponding to the loss of $CO_2$ from sn2 $RCOO^-$ was also observed with m/z 259.2. Data processing of fragment ions spectra was carried out with Biovia Draw 19,1© and MassLynx 4,1© software with the help of Lipid Maps Lipidomics Gatway®.

Statistical analysis was performed using two-way ANOVA, completed by Bonferroni post-tests to compare means variations between the two groups of animals for each PE(P) subspecies. No significant differences between CTL and HFHF were observed whatever the organ considered.

#### Supplementary Figure 8: PI Double-Bond (DB) index and DHA to AA ratios in various organs as a function of the diet.

Total lipids were extracted and phospholipid species were purified and analyzed by ESI-MS from samples corresponding to the indicated organs obtained either rats fed with a normal (CTL) or HFHF diet (HFHF), as described in the "Materials and Methods" section. The relative percentage of saturated (DB=0: no double bonds) versus monounsaturated (DB=1: one double bond), diunsaturated (DB=2: two double bonds) and polyunsaturated (DB > 2: > two double bonds) phosphatidylinositol (PI) species was obtained from the PI subspecies distribution displayed in Supplementary Fig. 4. The ratio of Arachidonic Acid (AA)- to Docosahexaenoic Acid (DHA)-containing PI subspecies in the various organs is also displayed.

#### Supplementary Figure 9: PE Double-Bond (DB) index and DHA to AA ratios in various organs as a function of the diet.

Total lipids were extracted and phospholipid species were purified and analyzed by ESI-MS from samples corresponding to the indicated organs obtained either rats fed with a normal (CTL) or HFHF diet (HFHF), as described in the "Materials and Methods" section. The relative percentage of saturated (DB=0: no double bonds) versus monounsaturated (DB=1: one double bond), diunsaturated (DB=2: two double bonds) and polyunsaturated (DB > 2: > two double bonds) phosphatidylethanolamine (PE) species was obtained from the PE subspecies distribution displayed in Supplementary Fig. 5. The ratio of Arachidonic Acid (AA)- to Docosahexaenoic Acid (DHA)-containing PE subspecies in the various organs is also displayed.

#### Supplementary Figure 10: PS Double-Bond (DB) index and DHA to AA ratios in various organs as a function of the diet.

Total lipids were extracted and phospholipid species were purified and analyzed by ESI-MS from samples corresponding to the indicated organs obtained either rats fed with a normal (CTL) or HFHF diet (HFHF), as described in the "Materials and Methods" section. The relative percentage of saturated (DB=0: no double bonds) versus monounsaturated (DB=1: one double bond), diunsaturated (DB=2: two double bonds) and polyunsaturated (DB > 2: > two double bonds) phosphatidylserine (PS) species was obtained from the PS subspecies distribution displayed in Supplementary Fig. 6. The ratio of Arachidonic Acid (AA)- to Docosahexaenoic Acid (DHA)-containing PS subspecies in the various organs is also displayed.

#### Supplementary Figure 11: PE(P) Double-Bond (DB) index and DHA to AA ratios in various organs as a function of the diet.

Total lipids were extracted and phospholipid species were purified and analyzed by ESI-MS from samples corresponding to the indicated organs obtained either rats fed with a normal (CTL) or HFHF diet (HFHF), as described in the "Materials and Methods" section. The relative percentage of saturated (DB=0: no double bonds) versus monounsaturated (DB=1: one double bond), diunsaturated (DB=2: two double bonds) and polyunsaturated (DB > 2: > two double bonds) PE(P) species was obtained from the PE(P) subspecies distribution displayed in Supplementary Fig. 7. The ratio of Arachidonic Acid (AA)-to Docosahexaenoic Acid (DHA)-containing PE(P) subspecies in the various organs is also displayed.

#### Supplementary Figure 12: Representative positive ion mass spectra of non-purified lipid extracts obtained from the liver and soleus of rats fed under a Normal (CTL) or HFHF diet.

Non-purified lipid extracts were analyzed in the positive ion mode with a scan range from 300 to 1200 m/z to detect Cholesterol, Diglycerides (DG), Triglycerides (TG) and PC species, as described in the "Materials and Methods" section. As shown, if the HFHF diet mainly induced variations in the relative proportions of PC species in the soleus, this diet also resulted in the deposition of neutral lipids in the liver (namely Cholesterol, and DG detected as their dehydrated forms, and TG) and the formation of new PC subspecies that were not detected in this organ under the CTL diet. The main PC subspecies identified by further fragmentation studies (MS/MS) are indicated in each case. Identified DG and TG subspecies corresponded to DG(16:0/18:1) (DG(34:1)), DG(18:1/18:1) (DG(36:2)), TG(16:0/18:1/18:1) (TG(52:2)) and TG(16:0/18:1/20:4) (TG54:5)).

#### Supplementary Figure 13: Neutral lipids in the liver and sole us muscle.

Total lipids were extracted from 30 mg of liver or soleus from either CTL or HFHF rats, as described under the "Materials and Methods" section. Neutral lipids were then analyzed as already described elsewhere (Ferreira et al., 2004). In brief, the final organic phase obtained after lipid extraction (Materials and Methods) was evaporated and lipids were dissolved in 100 $\mu$l of hexane. Neutral lipids (Triglycerides (TG), Diglycerides (DG), Sterylesters (SE) and Nonesterified Fatty Acids (NEFA)) were separated by Thin Layer Chromatography in hexane/diethyl ether/acetic acid (80:20:1, by vol.) on precoated Silica gel 60 F254 plates (Merck). Lipids were visualized under UV after vaporization of a primuline solution (0.05 mg/ml in acetone/water (80:20, v/v)) and identified by using appropriate standards. Representative results obtained from two rats (1 and 2) fed either with a normal (CTL) or a HFHF diet (HFHF) are displayed.

#### Supplementary Figure 14: Langendorff protocol

A: After the hearts have been set-up according to the Langendorff method, stabilization of contractile parameters was assessed and the hearts were submitted to 3 successive experimental phases of 30 min each: stabilization, global ischemia and reperfusion. B: Typical recording of left ventricular pressure obtained during the protocol described in A.

#### Supplementary Figure 15: Ischemia

Ischemic (upper) and reperfusion (lower) parameters were recorded in control (n=11) and HFHF (n=12) rats. Values from the two experimental groups were compared using repeated measures analysis of variance followed by Bonferroni post-hoc comparisons. A p value of less than 0.05 was considered as statistically significant.

#### Supplementary Table 1: Percentage of the various fatty acids in the HFHF diet

The percentage of the various fatty acids in the standard (CTL; MUCEDOLA, Settimo Milanese, Italy) and “WESTERN RD” (SDS, Special Diets Services, Saint Gratien, France) are as provided by the supplier. The name, the nature (saturated or unsaturated), the total carbon chain length (x) and number of carbon-carbon double bounds (y) are indicated for each fatty acid. N. D.: Not Detectable.

| | | | Standard (CTL) | Western RD |
|---|---|---|---|---|
| | **Name** | **x:y** | **% in the diet (m/m)** | **% in the diet (m/m)** |
| **Saturated Fatty Acids** | Lauric acid | 12:0 | 0.006 | 0.62 |
| | Myristic acid | 14:0 | 0.003 | 2.00 |
| | Palmitic acid | 16:0 | 0.383 | 6.07 |
| | Stearic acid | 18:0 | 0.084 | 2.66 |
| **Unsaturated Fatty Acids** | Myristoleic acid | 14:1 | N. D. | 0.17 |
| | Palmitoleic acid | 16:1 | 0.024 | 0.38 |
| | Oleic Acid | 18:1 | 0.511 | 5.64 |
| | Linoleic acid (ω-6) | 18:2 | 1.298 | 0.83 |
| | Linolenic acid (ω-3) | 18:3 | 0.247 | 0.13 |

#### Supplementary Reference

Ferreira, T., Regnacq, M., Alimardani, P., Moreau-Vauzelle, C., and Berges, T. (2004). Lipid dynamics in yeast under haem-induced unsaturated fatty acid and / or sterol depletion. Biochem J *378*, 899-908.
