## Supplementary Figures for "A comprehensive study of Phospholipid fatty acid rearrangements in the early onset of the metabolic syndrome: correlations to organ dysfunction"

### Supplementary Fig. 1

A

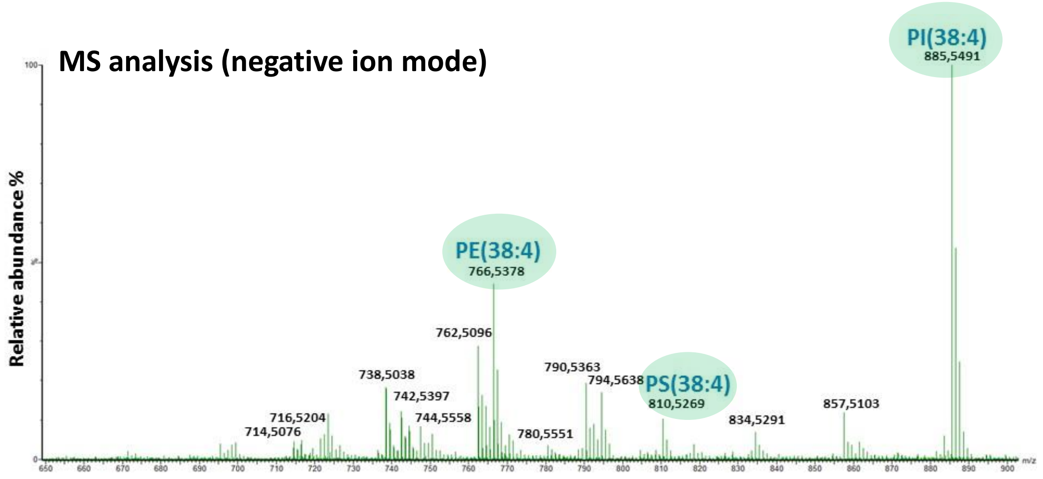


B

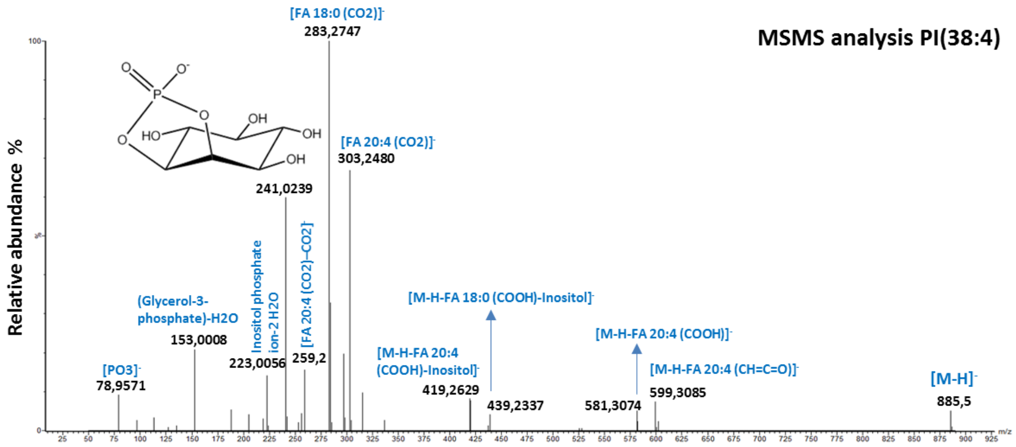


C

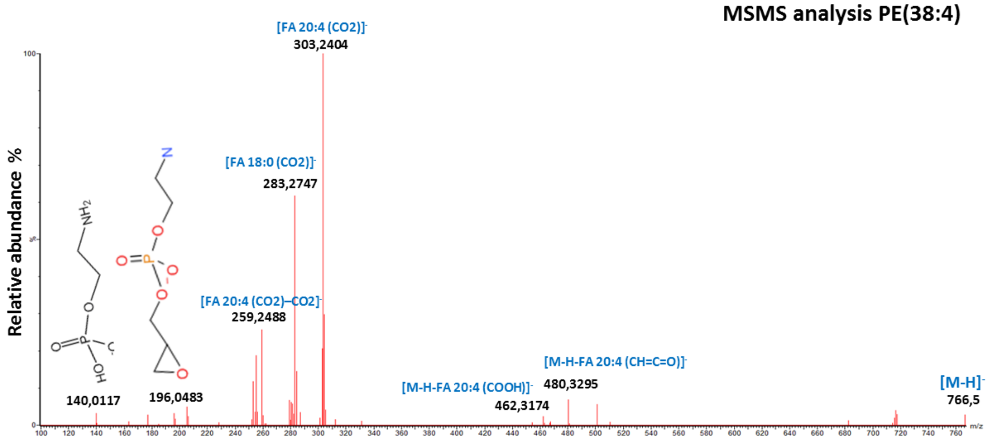


D

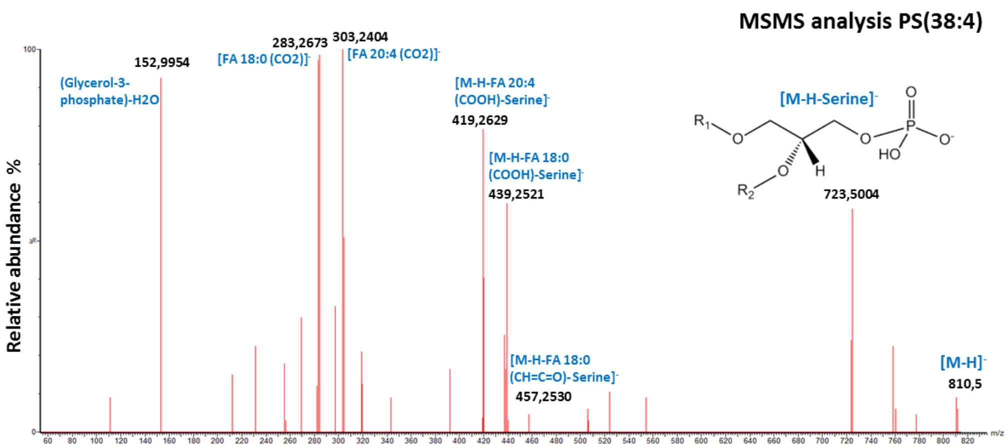

### Supplementary Fig. 2

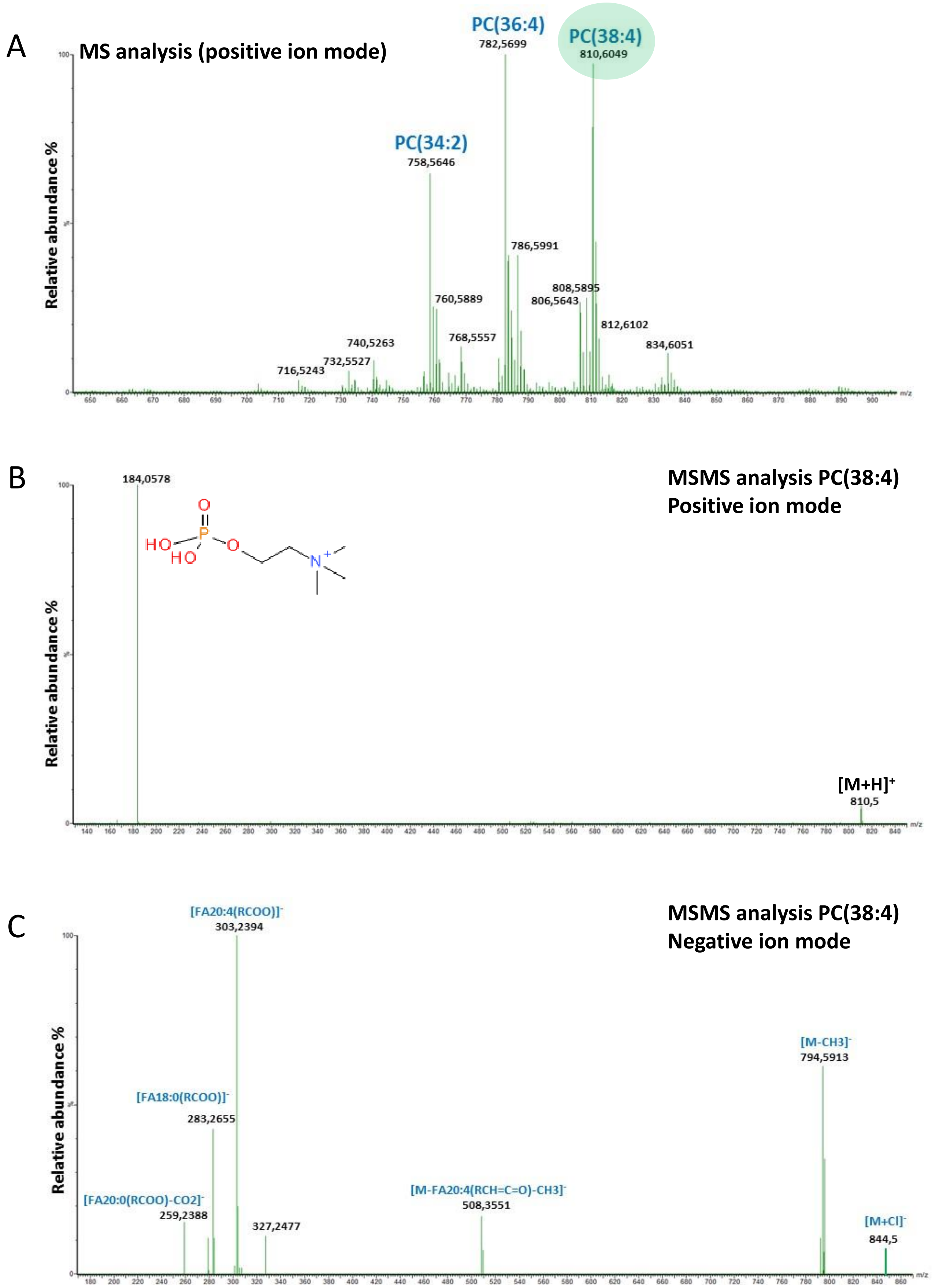

### Supplementary Fig. 3

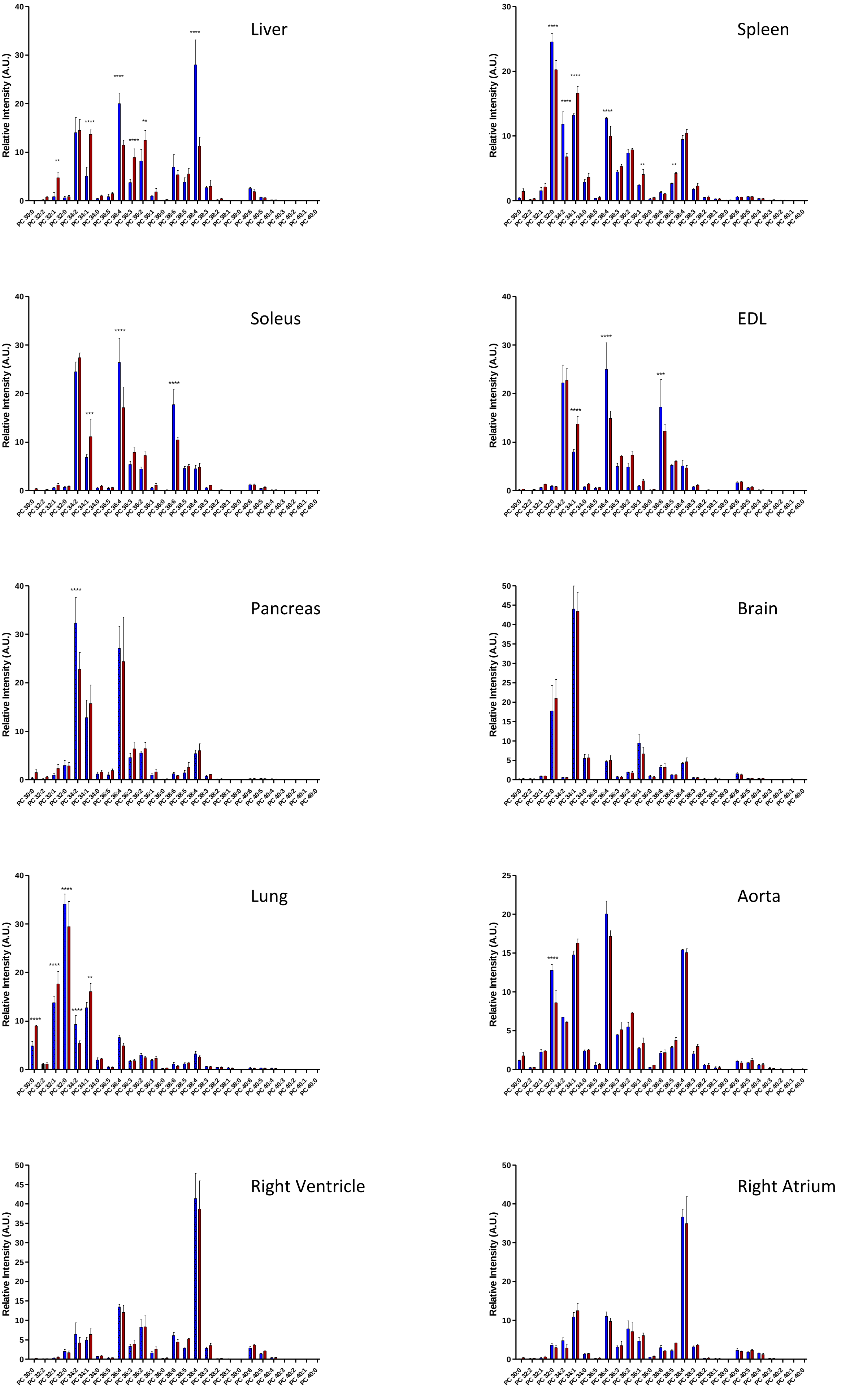

### Supplementary Fig. 4

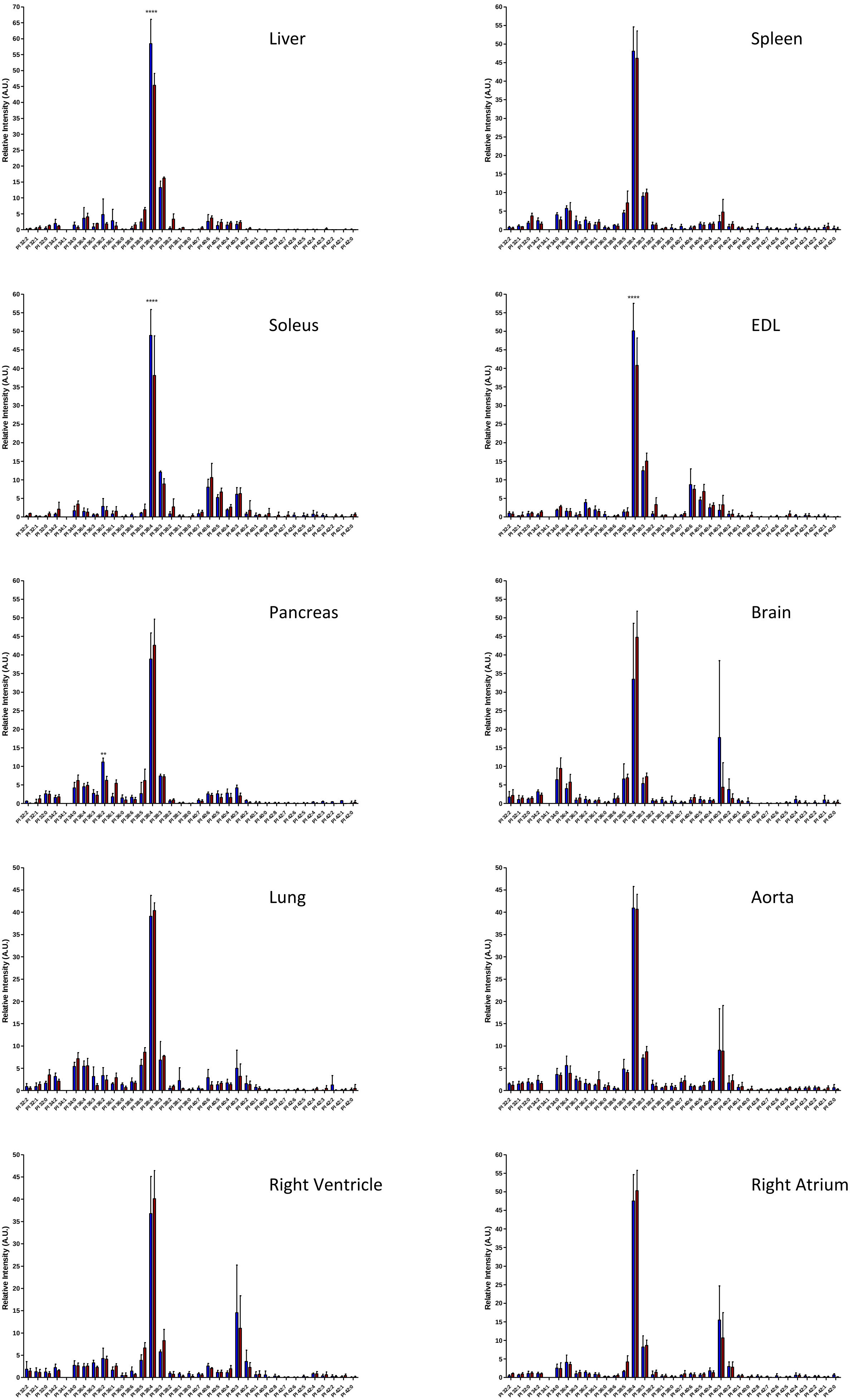

### Supplementary Fig. 5

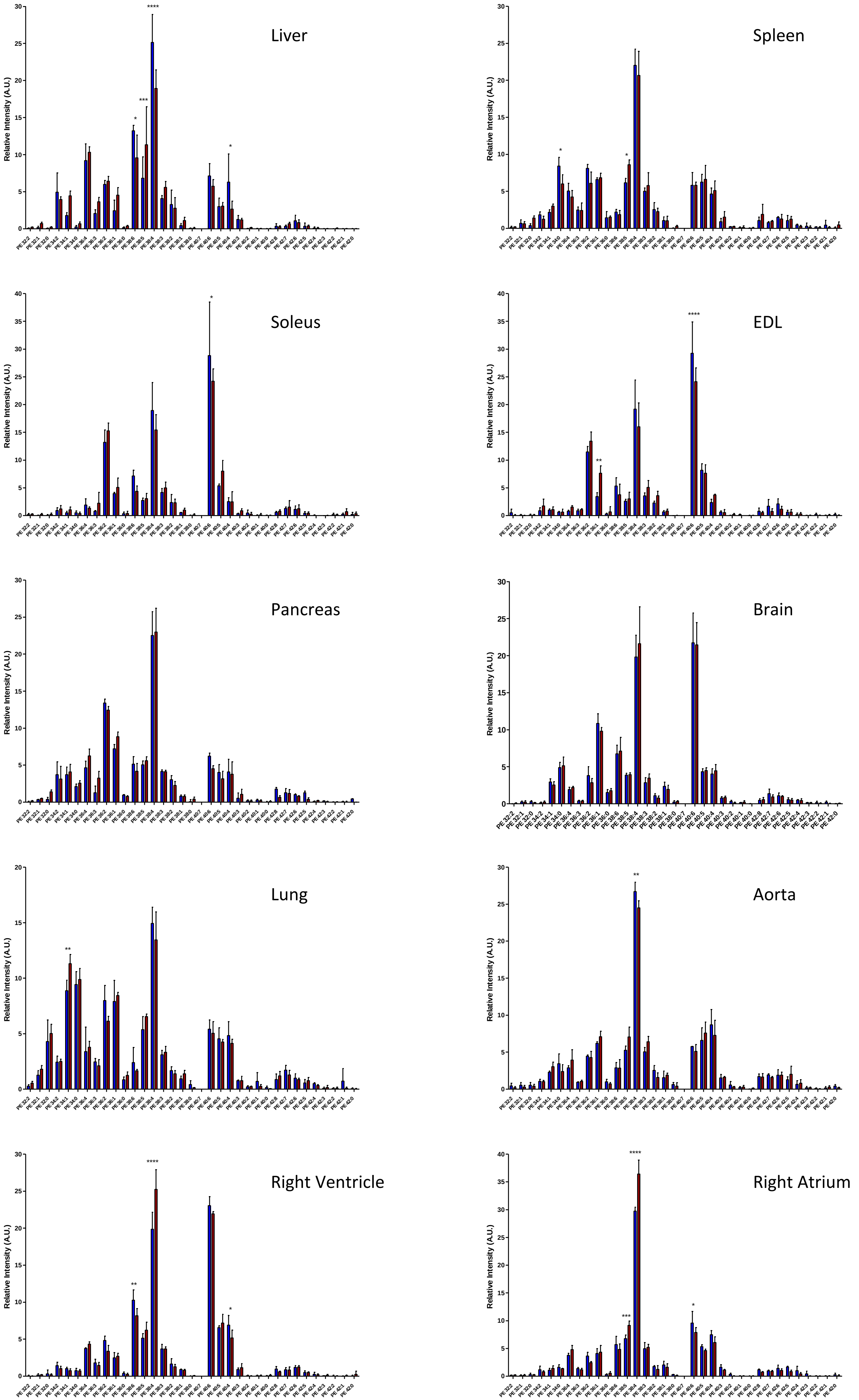

### Supplementary Fig. 6

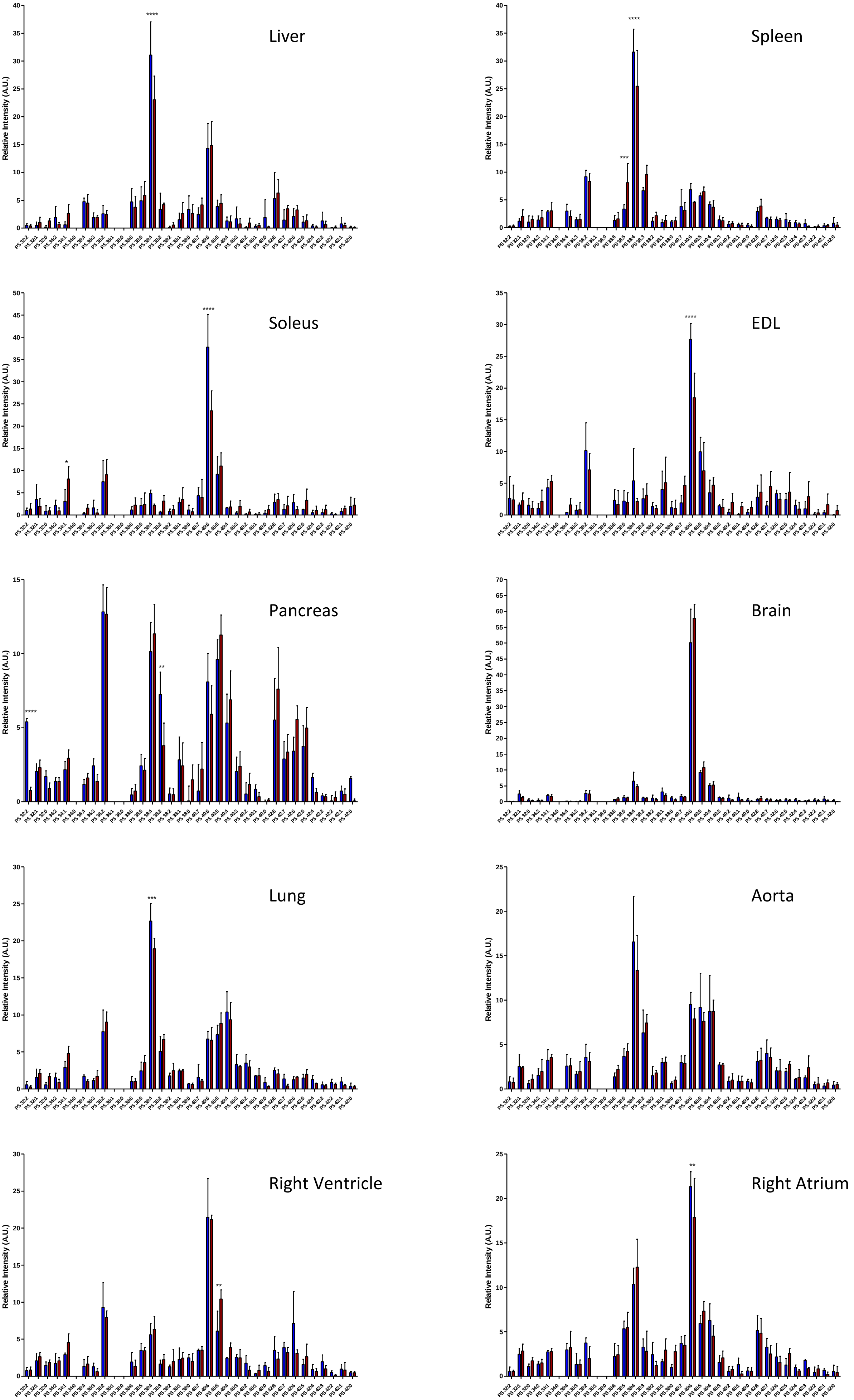

### Supplementary Fig. 7

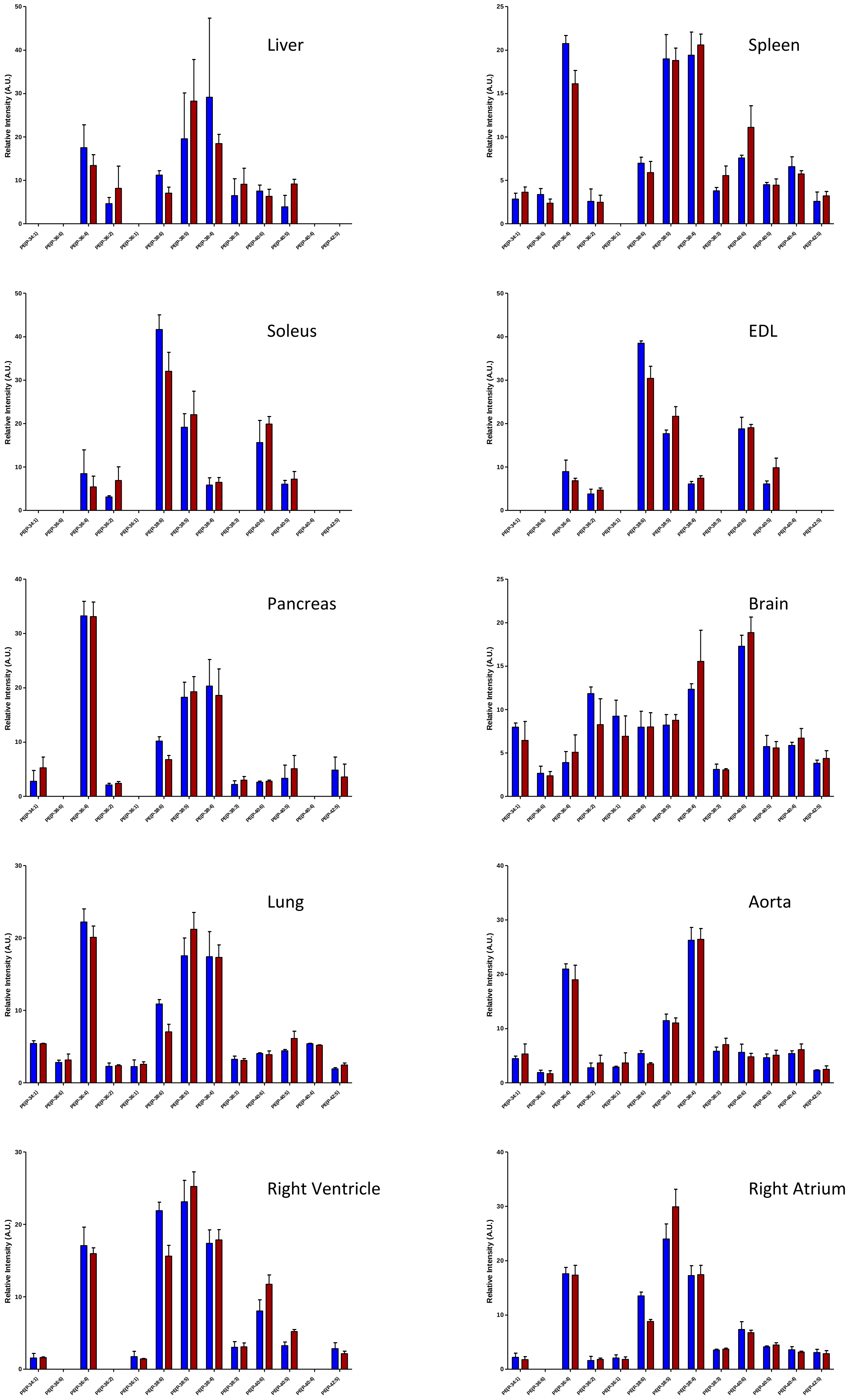

### Supplementary Fig. 8

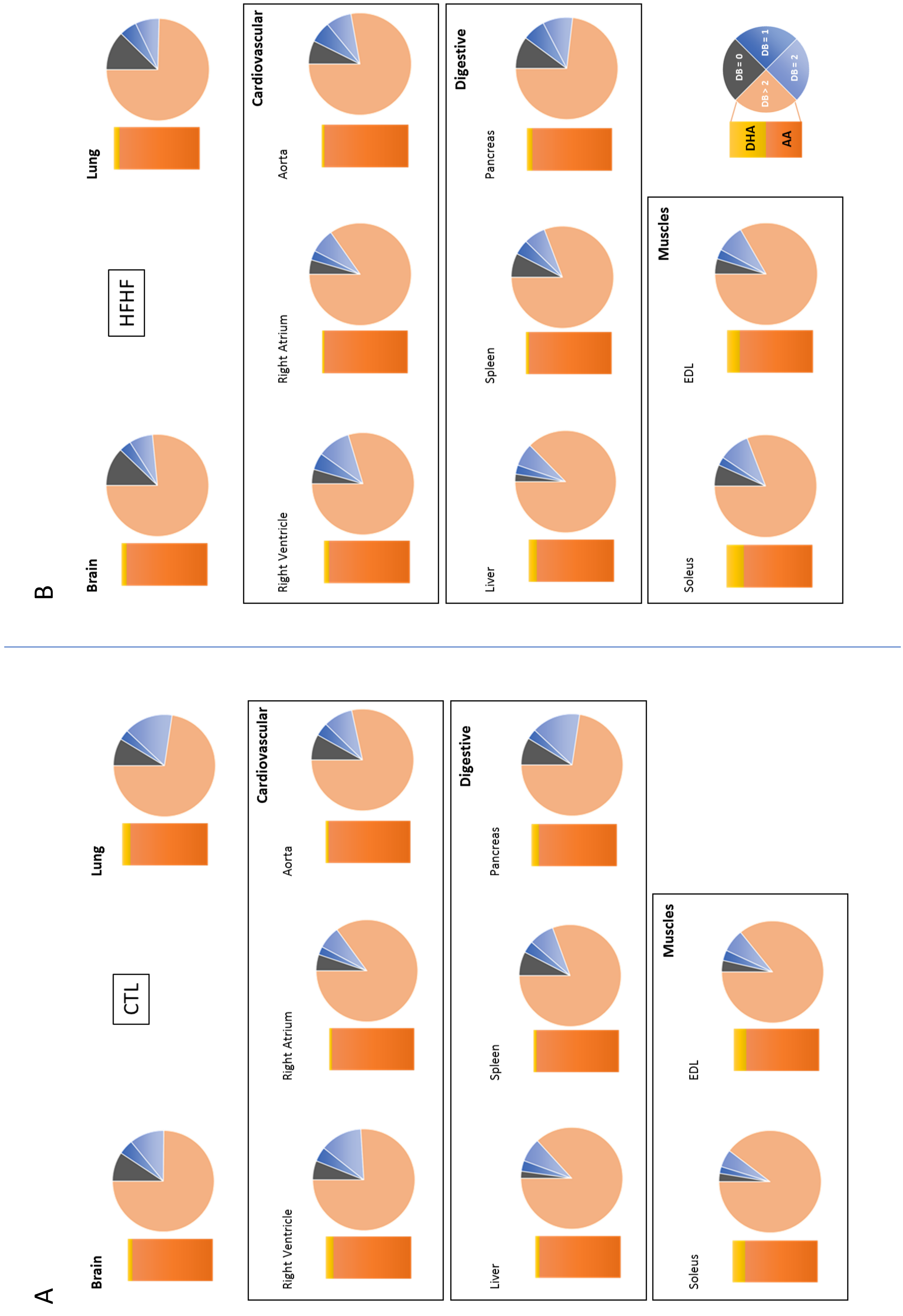

### Supplementary Fig. 9

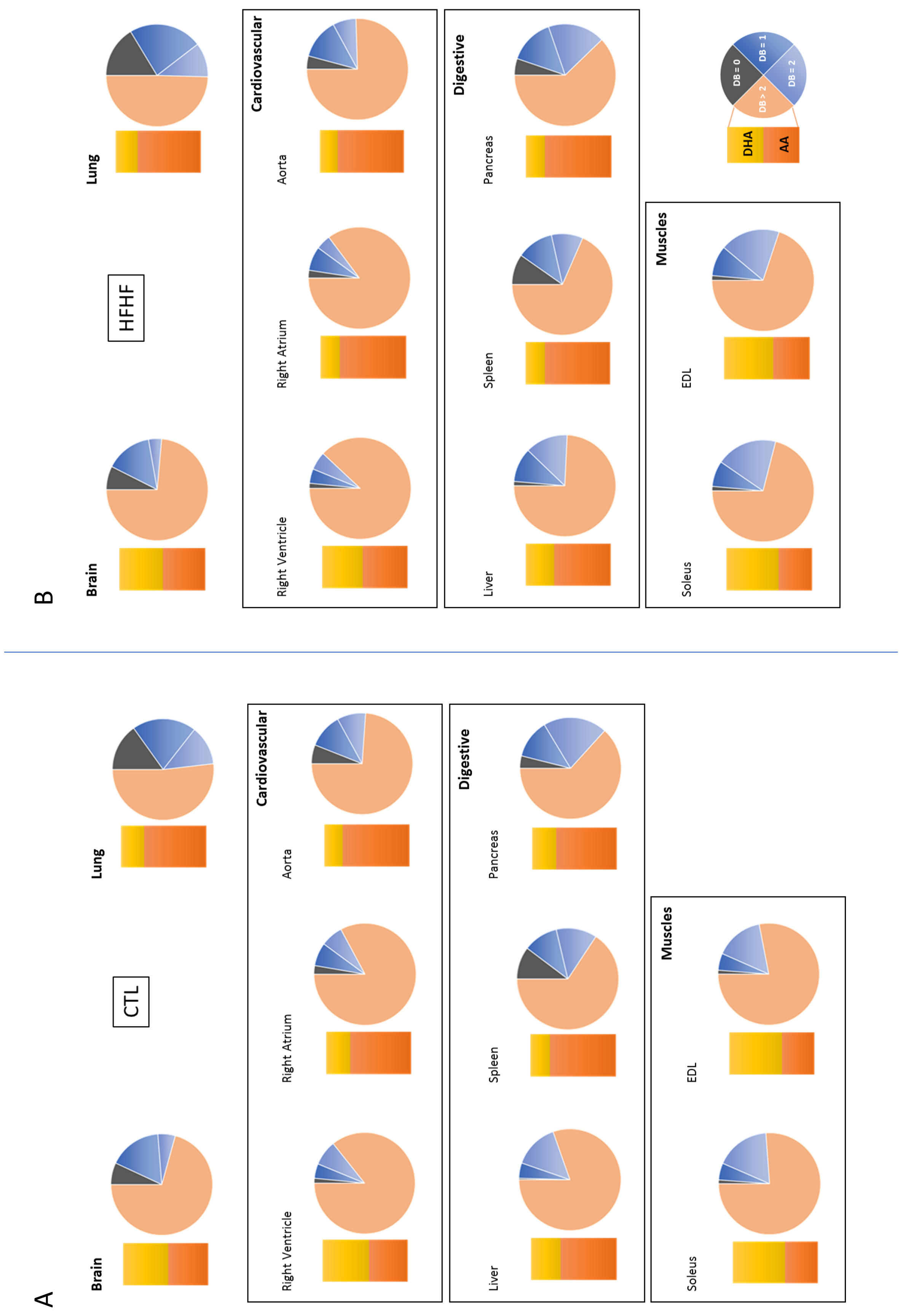

### Supplementary Fig. 10

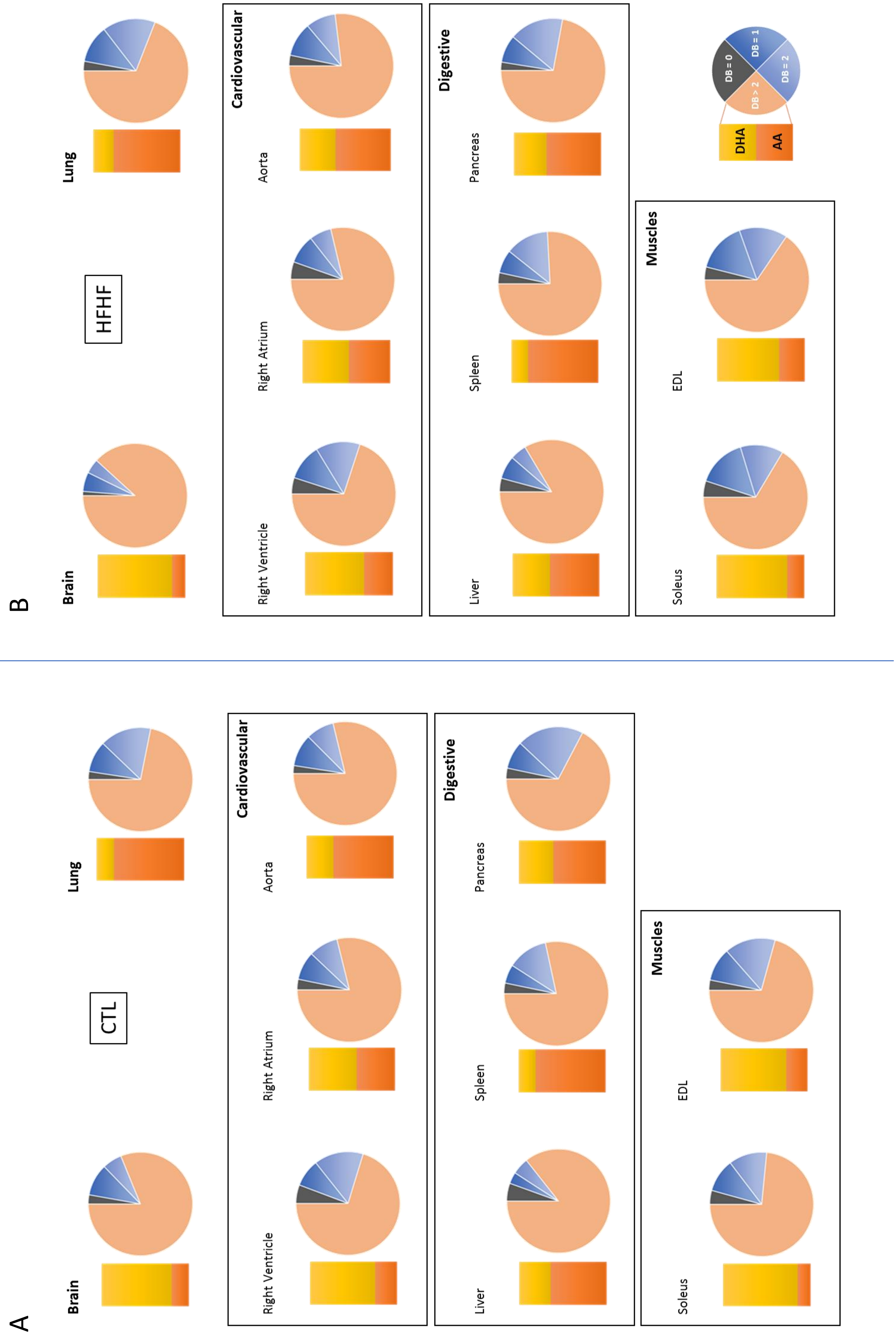

### Supplementary Fig. 11

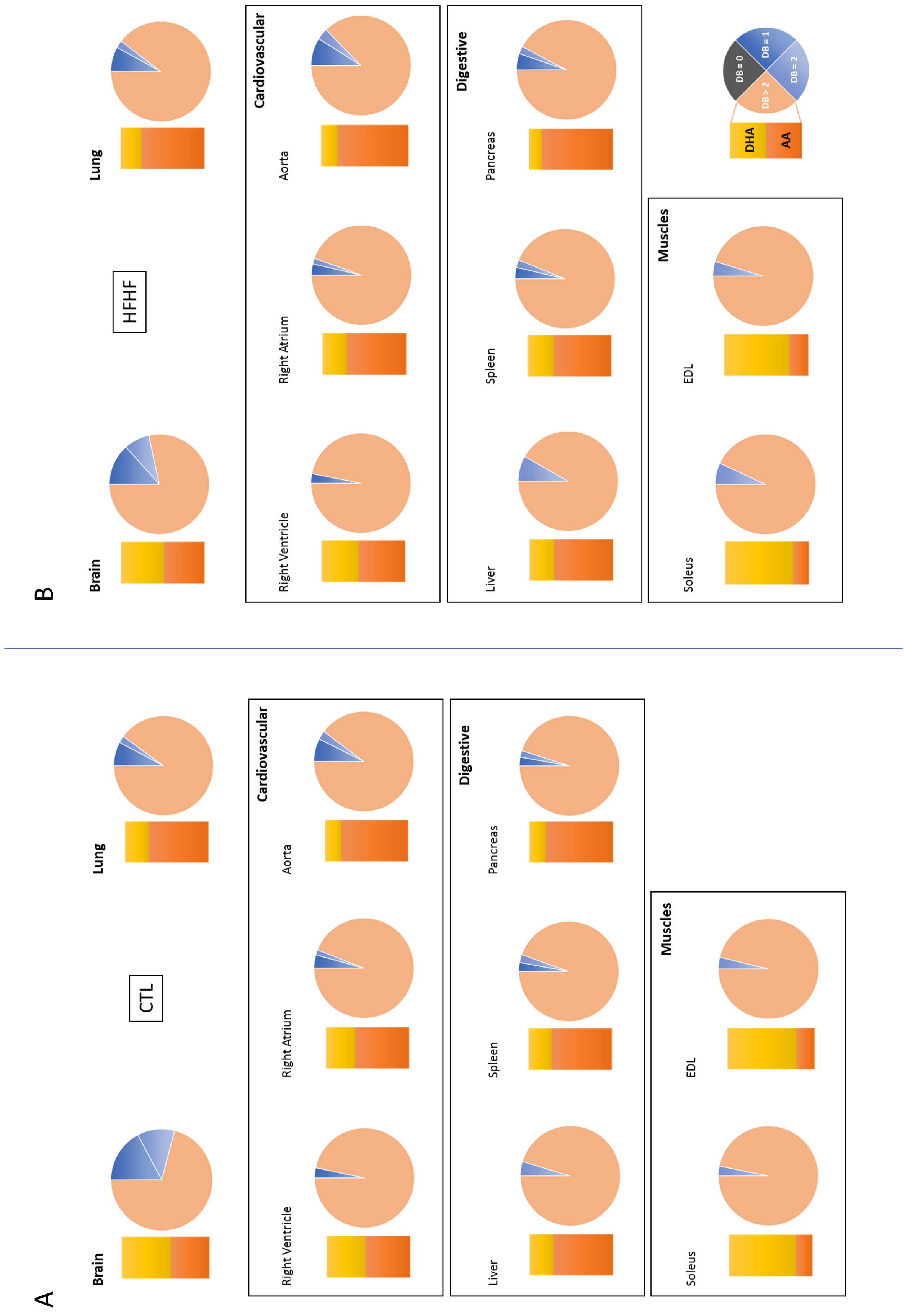

### Supplementary Fig. 12

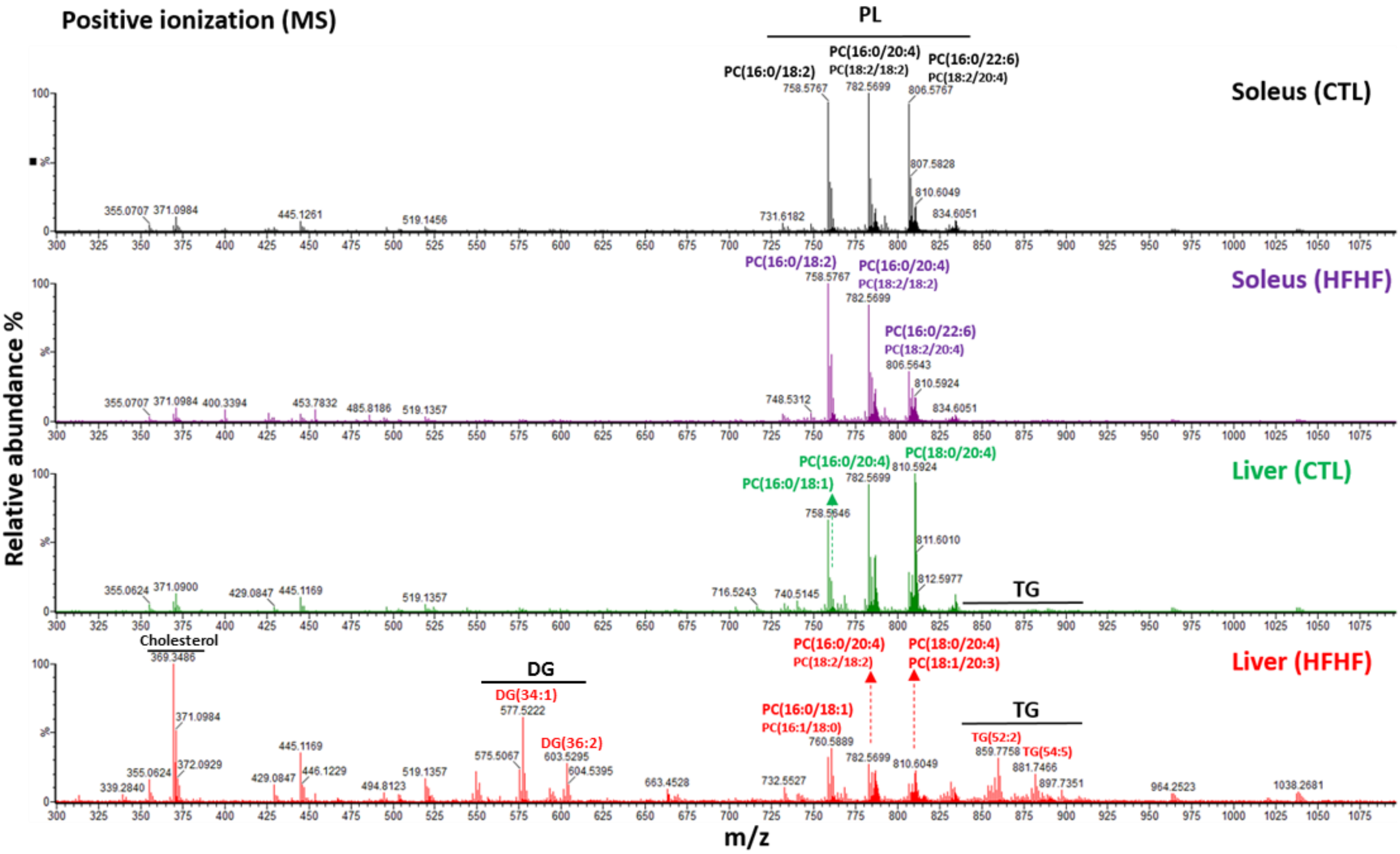

Supplementary Fig. 13

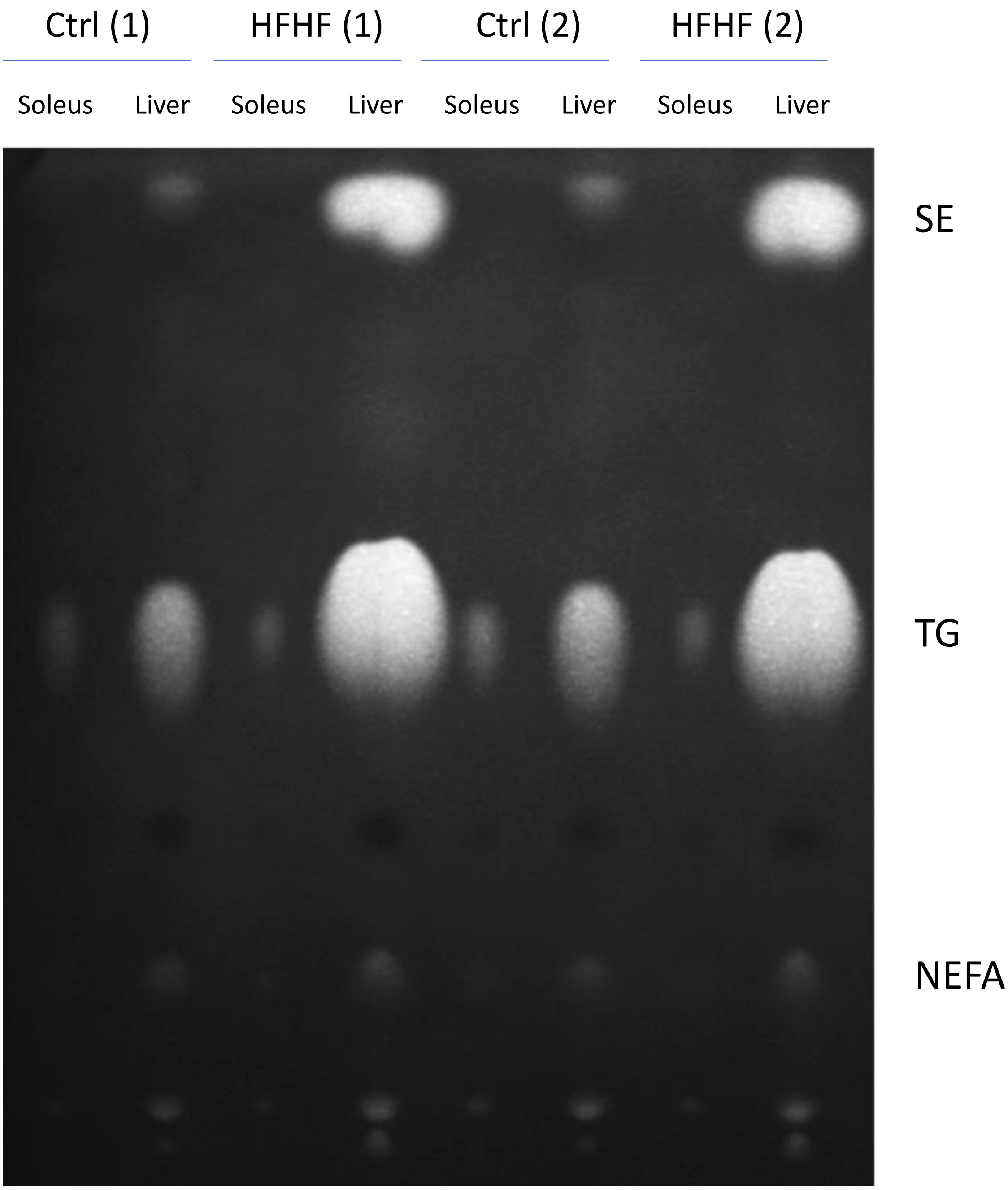

Supplementary Fig. 14

A

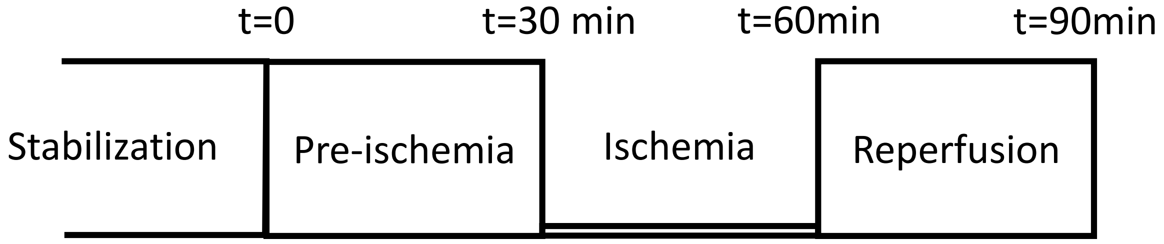


B

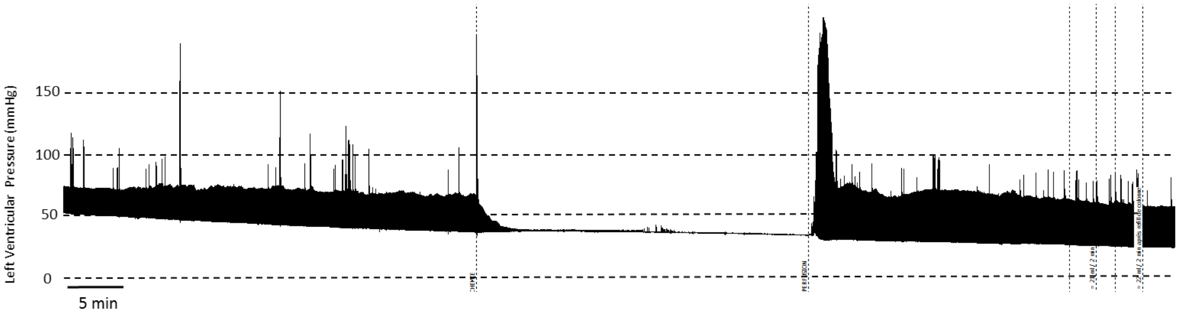

### Supplementary Fig. 15

#### ISCHEMIA

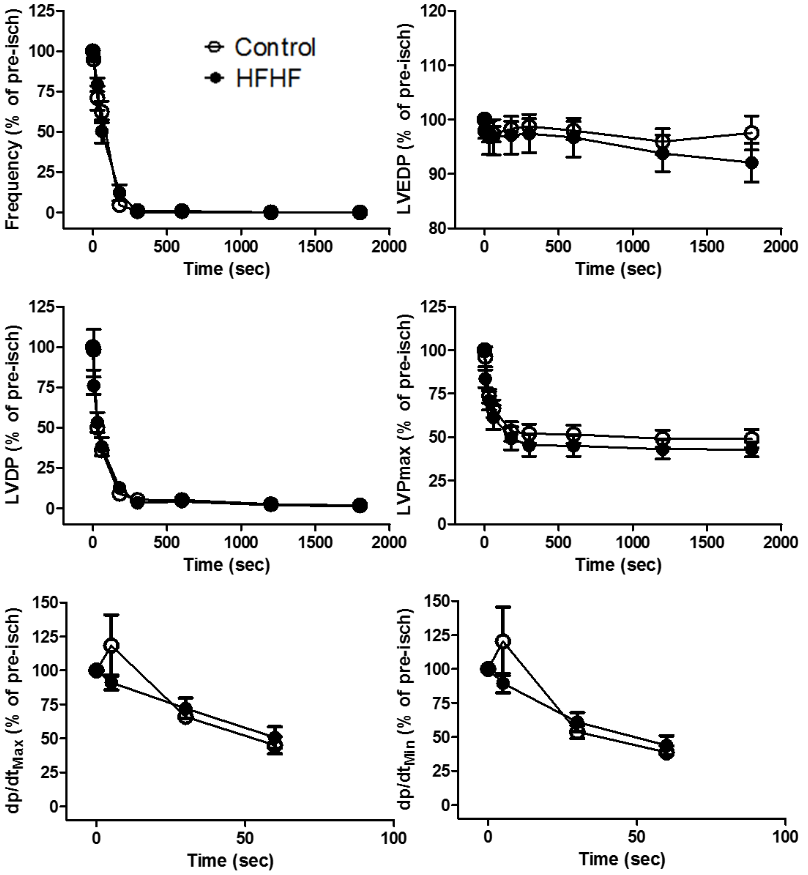


#### REPERFUSION

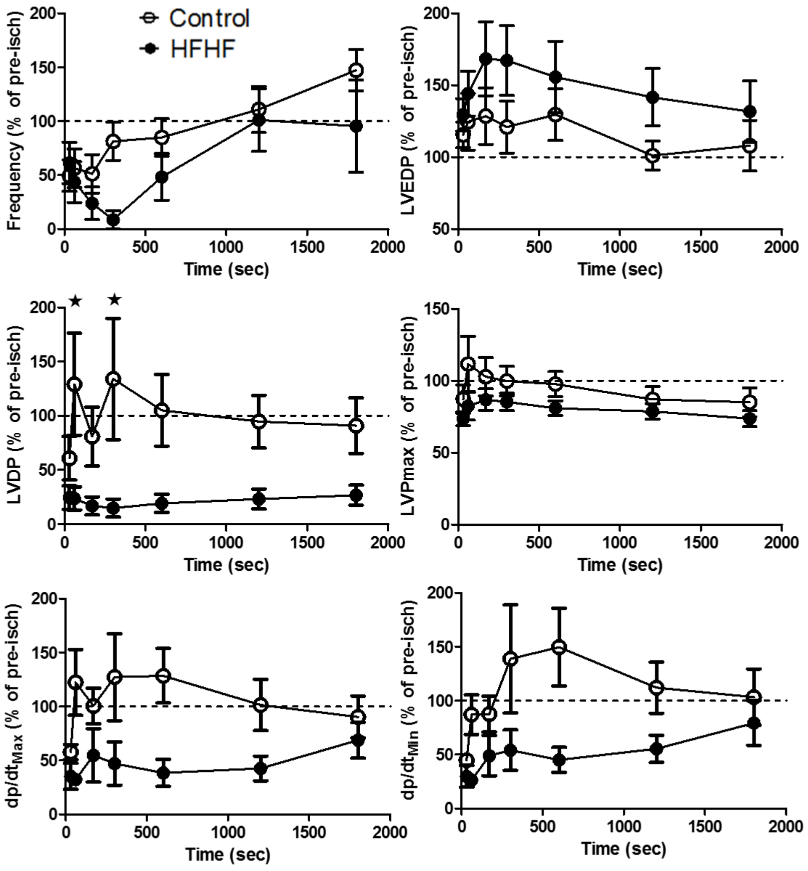
